# A CRISPR/*Sp*Cas9M-reporting system for efficient and rapid genome editing in *Caulobacter crescentus*

**DOI:** 10.1101/2024.12.02.626314

**Authors:** Jingxian Sun, Xin Yu, Guiyue Tang, Mengqing Chen, Yixin Zheng, Yucan Hu, Qingmei Li, Xiaoyang Li, Ningning Li, Zhongyue Li, Yingying Li, Ning Lu, Wei Tan, Yujiao Yang, Xiaoye Zeng, Guohong Zhao, Hailong Wang, Lei Dai, Guo-Ping Zhao, Lianzhong Ai, Wei Zhao

## Abstract

As members of the α-proteobacteria group, *Caulobacter crescentus* and its relatives are known for their asymmetric life cycle and comprehensive applications in gene delivery, agricultural biotechnology, and the production of high-value compounds. However, genetic manipulations of these bacteria are often time-consuming and labor-intensive due to the lack of efficient genome editing tools. Here, we report a practical CRISPR/*Sp*Cas9M-reporting system that overcomes the limitations of *Sp*Cas9 expression, enabling efficient, markerless, and rapid genome editing in *C. crescentus*. As a demonstration, we successfully knocked out two genes encoding the scaffold proteins, achieving apparent editing efficiencies up to 80%. Key components, including the Cas protein, Cas inducer, sgRNA, homologous arms, and reporter, were systematically analyzed and optimized to enhance the editing efficiency or decrease the cell lethality. A nearly zero off-target ratio was observed after the curing of the editor plasmid in editing strains. Furthermore, we applied the CRISPR/*Sp*Cas9M-reporting system to two *C. crescentus* relatives, *Agrobacterium fabrum* and *Sinorhizobium meliloti*, establishing it as an efficient and reliable editing strategy. We anticipate that this system could be applied to other hard-to-edit organisms, accelerating both basic and applied research in α-proteobacteria.

## Introduction

As one of the largest groups of bacteria, α-proteobacteria are widespread across diverse ecological niches but share a common evolutionary ancestor. This group encompasses a wide array of microorganisms, including phototrophs (e.g., *Rhodobacter*), plant symbionts (e.g., *Sinorhizobium*), animal pathogens (e.g., *Brucella*), and several genera that metabolize C1 compounds (e.g., *Methylobacterium*) (1–3). The α-proteobacteria are also the close relatives of proto-mitochondrion, the latter of which is considered as the one of origins that generated aerobic eukaryotes (4, 5).

*Caulobacter crescentus* belongs to Caulobacterales of α-proteobacteria, and serves as a model organism for studying asymmetric cell division (*SI Appendix*, Fig. S1) (6). Unlike many other α-proteobacteria, *C. crescentus* is a free-living bacterium commonly found in oligotrophic freshwater environments, such as lakes and rivers (7, 8). In each cell cycle, *C. crescentus* divides into two distinct daughter cells: a motile swarmer cell and a stationary stalked cell (9). This asymmetric division, a trait shared by many α-proteobacteria and virtually all eukaryotes, makes *C. crescentus* an excellent model for investigating cell differentiation and cell cycle regulation. As a plant pathogen or symbiont, *Agrobacterium tumefaciens* and *Sinorhizobium meliloti* belong to Rhizobiales of α-proteobacteria and also exhibit asymmetric division (10–12). *A. tumefaciens* has long been used in plant genome engineering through delivering sequences using the T-DNA binary vectors (13, 14), while *S. meliloti* is known for its nitrogen-fixing capacity with legumes, which has agricultural significance (15, 16).

Despite their importance in both basic research and industrial/agricultural applications, genetic manipulation of *C. crescentus* and its relatives remains time-consuming and labor-intensive. The current genetic tool used for *C. crescentus* genome editing was first developed in 1991 by Ely et al., using the *sacB* counterselection (17). However, this method often requires more than three weeks to complete a single editing cycle and exhibits relatively low editing efficiency (18). In 2006, Thanbichler et al. reported a single-crossover method that facilitates efficient gene integration in *C. crescentus* (18, 19). Nevertheless, this method introduces the whole editor plasmid with selection markers into the chromosome and is unable to facilitate gene deletion. More recently, a novel tool for gene silencing was developed using the CRISPR (clustered regularly interspaced short palindromic repeats) interference system (20); however, it remains difficult to achieve gene deletion or insertion with this method.

Scientists have attempted to develop efficient genome-editing tools for *C. crescentus* based on homologous recombination (HR), but these efforts have largely been unsuccessful (17, 18). Similar challenges exist for *S. meliloti* and *A. tumefaciens*, two evolutionary relatives of *C. crescentus* within α-proteobacteria (Fig. S1). It has been suggested that all these bacteria possess very weak cellular recombinase activity, though no direct evidence has been provided (21, 22). The lack of efficient, markless, and rapid genome editing methods has significantly hindered functional gene analysis and rational genome engineering in this group of organisms, highlighting an urgent need for new genetic tools.

To address these challenges, we developed a CRISPR/Cas-assisted HR method for *C. crescentus* using an all-in-one editor plasmid. To achieve efficient genome editing, we screened various Cas proteins and systematically optimized the editor plasmid, particularly introduced a CRISPR/*Sp*Cas9M-reporting system. The reporting system facilitates the identification of CRISPR escape events and substantially increases the apparent editing efficiency in *C. crescentus*. We successfully applied this design to two relatives of *C. crescentus* (*A. tumefaciens* and *S. meliloti*), demonstrating it as a general and efficient genome editing strategy. Our method enables targeted gene deletion and insertion within less than a week, producing in-frame and markerless edits on the chromosomes. We anticipate that this practical tool could be applicable to other difficult-to-edit organisms, which would significantly benefit both basic and applied research in α-proteobacteria.

## Results

### Construction and optimization of a HR-assisted CRISPR/Cas system in *C. crescentus*

We first constructed an HR-assisted CRISPR/Cas system using the *Streptococcus pyogenes* Cas9 (*Sp*Cas9), which is encoded on the well-characterized *C. crescentus* replicating plasmid pBXMCS2 (19). We used a vanillate-inducible promoter (P*^van^*) to drive *Sp*Cas9 and a constitutive promoter (P^J23119^) to transcribe the single guide RNA (sgRNA). We also designed a pair of homologous arms (H-arms) to repair DNA double-strand breaks (DSBs) caused by Cas cleavage. In order to achieve markerless editing, no antibiotic resistant gene was designed between the H-arms. After electroporating the editor plasmid into *C. crescentus* and selection on the presence of plasmid, however, no colony or very few colonies were obtained compared to the controls without sgRNA or with non-targeting sgRNA (*SI Appendix*, Fig. S2*A*). Moreover, the surviving colonies did not have any editing at the targeted sites. These results are consistent with previous observations in *A. tumefaciens* (21) and *S. meliloti* (22), indicating that the CRISPR/*Sp*Cas9 cleavage is lethal and very weak HR activity exist in *C. crescentus* cells.

In order to achieve effective targeted genome editing, we systematically analyzed our editor plasmid and optimized it in three key domains. First, we screened the Cas proteins from diverse organisms, including *Sp*Cas9, *Francisella novicida* Cas12a (*Fn*Cas12a), *Streptococcus thermophilus* CRISPR1-Cas9 (*Sth*1Cas9), and *Streptococcus thermophilus* CRISPR3-Cas9 (*Sth*3Cas9), to test their editing efficiencies in *C. crescentus*. Two genes encoding for the scaffold proteins SpmX and PodJ were selected as the editing targets because they are known as nonessential genes in *C. crescentus* (23). No edits were observed for all the Cas proteins mentioned above. However, a small group (∼15%) of edited colonies were detected after we optimized the *Sp*Cas9 coding sequence (designated as *Sp*Cas9M), using the codon preference of *C. crescentus* (Fig. 1*A*, 1*B* and *SI Appendix*, Table S1). No obvious edits were detected for the other optimized Cas proteins. This result suggests the expression of *Sp*Cas9 is related to the editing efficiency in *C. crescentus*.

**Fig. 1.**
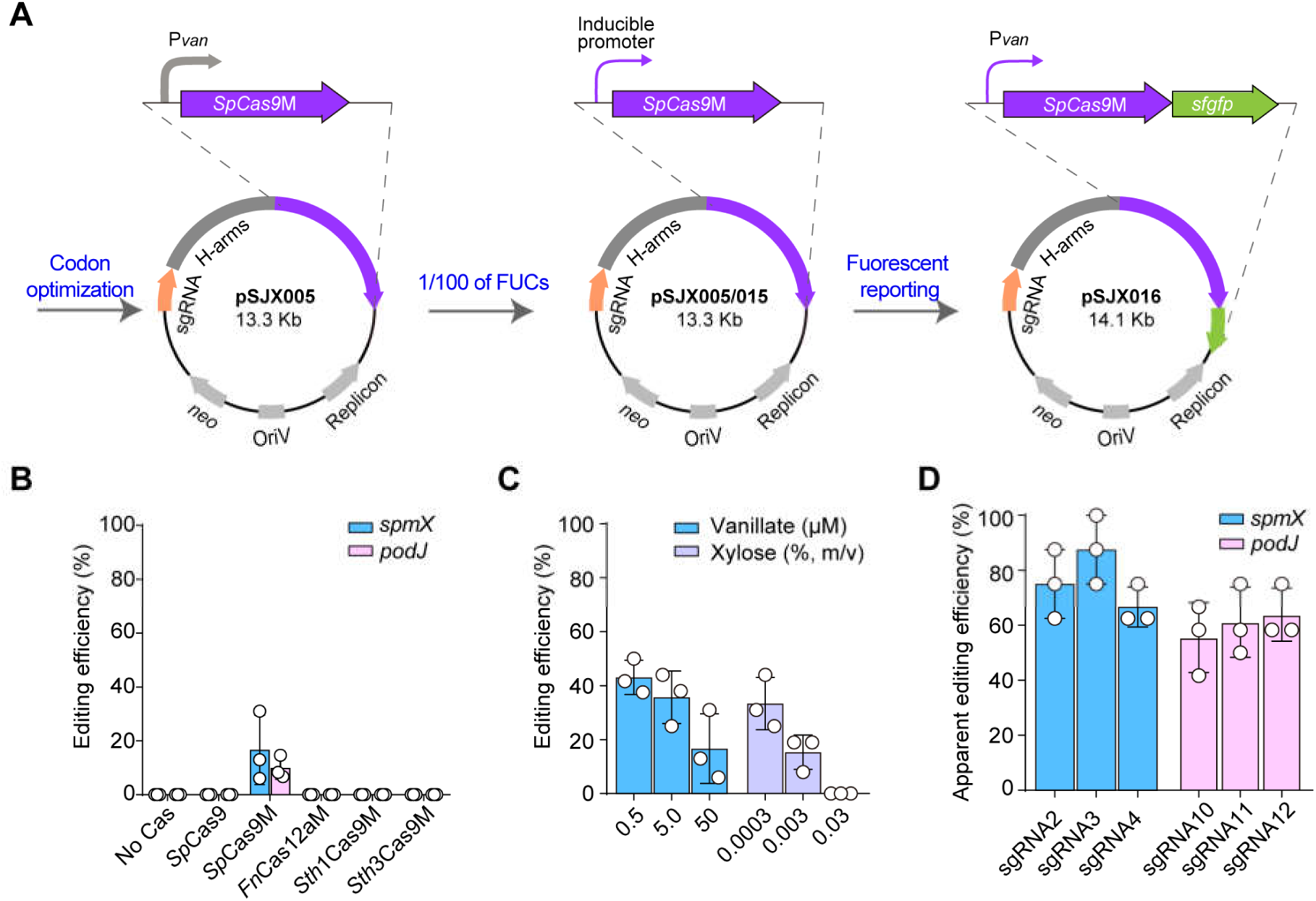
Development of a CRISPR/*Sp*Cas9M-reporting system in *C. crescentus*. (*A*) A flow diagram shows that three key optimization steps were used to enhance the genome editing efficiency in *C. crescentus*, including codon optimization of the *cas* gene, a decrease of *Sp*Cas9M expression level, and the creation of an *Sp*Cas9M-sfGFP reporting system. (*B*)-(*D*) The genome editing efficiency (or apparent editing efficiency, AEE) increases gradually as the system optimization progressed. (*B*) A small edited group was observed after optimizing the codon of *Sp*Cas9. Two genes encoding for the scaffold proteins SpmX and PodJ were used as the editing targets. (*C*) Comparison of the editing efficiency using different inducer concentrations for P*^van^* and P*^xyl^*. The *spmX* gene was used as the editing target. (*D*) The apparent editing efficiency reaches ∼80% for *spmX* and ∼60% for *podJ* using sfGFP as the fluorescent reporter. Three sgRNAs were designed and tested for each target gene. The genome editing efficiency (or AEE) was calculated by dividing the edited clones by all the tested clones (*n* = 36). Data are means ± SEM from 3 independent biological replicates.

We then asked if we could improve the editing efficiency by manipulating the expression level of *Sp*Cas9M. Two inducible promoters(19), P*^van^* and the xylose-inducible promoter P*^xyl^* with different concentrations of inducers were tested to evaluate their influence on the editing of *spmX* in *C. crescentus*. Unexpectedly, the editing efficiency decreased with increasing concentrations of xylose or vanillate. The use of xylose in a frequently-<u>u</u>sed <u>c</u>oncentration (FUC: 0.03%, m/V) resulted in no edits, while the use of inducers at 100x less than the FUCs displayed editing efficiency up to 40% (Fig. 1*C*). These results indicate that the relatively low expression of *Sp*Cas9M may be more favorable to genome editing in *C. crescentus*. Consistent with this conclusion, a constitutive high expression of *Sp*Cas9M using P^J23119^ resulted no edited clones (data not shown).

We therefore use the inducers with the concentrations 100x less than the FUCs to induce *Sp*Cas9M. However, the editing efficiency was still not high, and particularly, was unstable for different targeted sites. When checked the editor plasmids in surviving colonies without edits (i.e., CRISPR escapers), we noticed significant deletions on all the coding sequence of *Sp*Cas9M, ranging from 545 bp to the whole *SpCas9*M sequence (*SI Appendix*, Fig. S2*B*). These observations indicate that possible anti-CRISPR systems may exist in *C. crescentus.* Nevertheless, we hypothesized that we could improve the apparent editing efficiency (AEE) by pre-identification and exclusion of these *Sp*Cas9M mutants. To do this, a superfolder GFP (sfGFP) protein was designed to fuse to the C-terminal of *Sp*Cas9M (Fig. 1*A*). Through the indication of fluorescent sfGFP, the clones abnormally expressing *Sp*Cas9M were identified and excluded before colony PCR. The AEEs reached 80% and 60% for SpmX and PodJ edits, respectively, after using the fluorescent reporter. The use of the sfGFP indicator was a demonstration of a practical genome editing method in *C. crescentus* (Fig. 1*C*). We named this method including the induce conditions as CRISPR/*Sp*Cas9M-reporting system.

### Streamlining the CRISPR/*Sp*Cas9M-reporting system

Based on the above results, we have developed an all-in-one editor plasmid pSJX016, which enabled efficient, markless, and rapid genome editing in *C. crescentus* (Fig. 2*A*). In this plasmid, the coding gene for the fused *Sp*Cas9M-sfGFP, the custom sgRNA, and the H-arms were assembled into a high-copy-number shuttle plasmid, pBXMCS2. When targeting a new specific site, the pSJX016 could be used as a start point by replacing the sgRNA and H-arms in just one Gibson assembly.

**Fig. 2.**
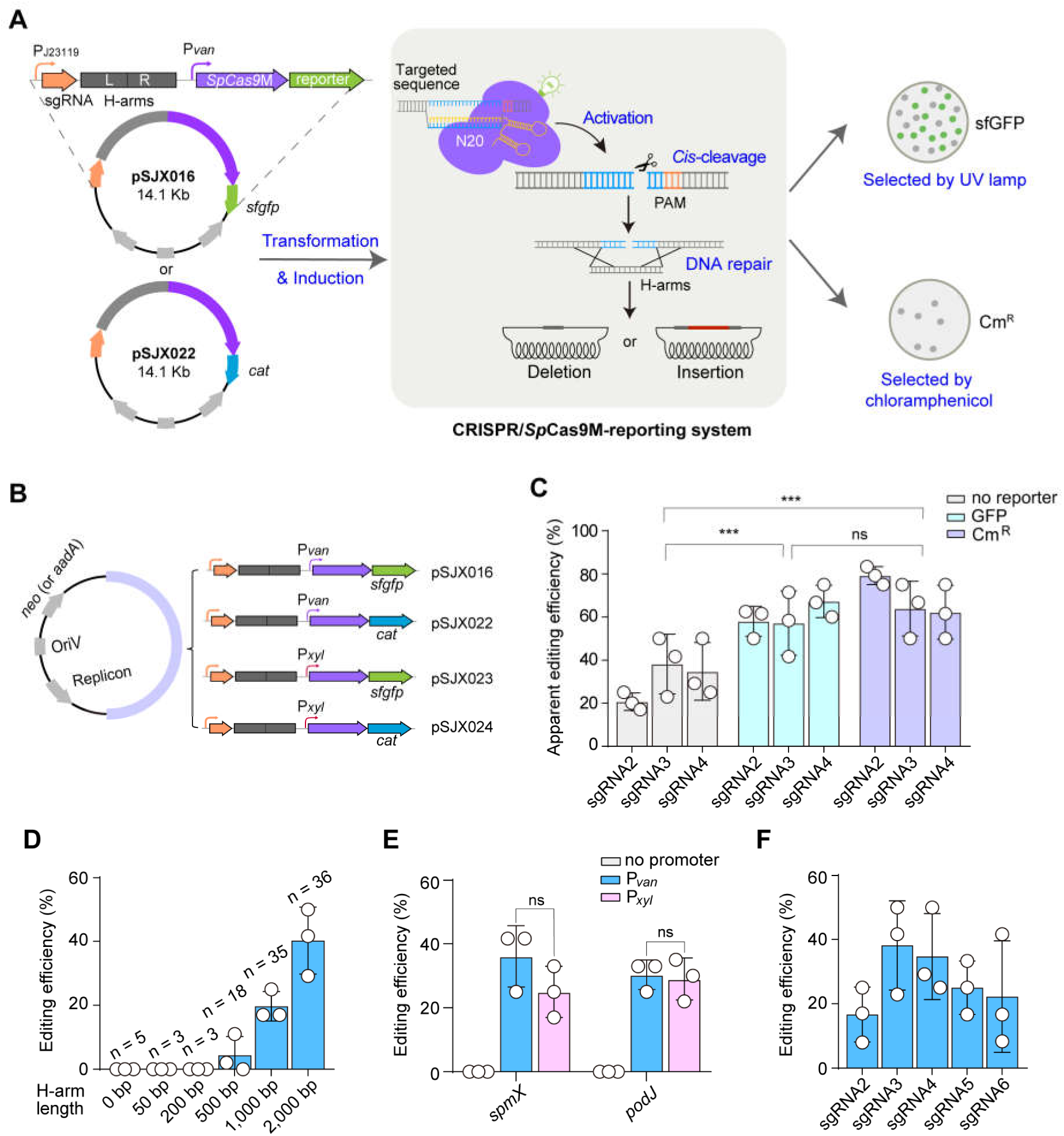
Characterization and streamlining of the CRISPR/*Sp*Cas9M editing system. (*A*) The all-in-one editor plasmid containing a P^J23119^ driven sgRNA, a P*^van^* driven *sp*Cas9M-sfGFP, and the corresponding H-arms was constructed and transformed into *C. crescentus.* The reporter gene could be expanded from the fluorescent *sfgfp* to the antibiotic resistance gene *cat*, which provided an alternative CRISPR/*Sp*Cas9M-reporting system. The clones in the reporting system could be pre-screening by using either blue light or chloramphenicol. (*B*) A representation of editor plasmids with different inducible promoters and reporter genes. (*C*) The CRISPR/*Sp*Cas9M-Cat system shows comparable AEEs to the fluorescent reporting system, but displays significantly higher AEEs than the non-reporting system. Three sgRNAs targeting the *podJ* gene were tested. The *p* value was determined by the unpaired *t* test. (*D*) The *spmX* editing efficiency increases gradually when the H-arms are elongated in the non-reporting system. The unilateral lengths of H-arms are shown on the X-axis. (*E*) No significant difference in editing efficiency between the inducible promoter P*^van^* and P*^xyl^*. Both *spmX* and *podJ* were used as the editing targets. Working concentrations of 0.5 µM vanillate and 0.0003% (m/V) xylose were used. The *p* value was determined by Welch’s paired *t*-test. (*F*) The editing efficiency varies between different sgRNAs. Five sgRNAs targeting *spmX* were designed. All data are presented as means ± SEM from 3 independent biological replicates. A total of 36 clones (*n*) were tested for each sample unless other indicated. ns, *p* > 0.05; ***, *p* ≤ 0.01.

A complete genome editing workflow using the CRISPR/*Sp*Cas9M-reporting system could be achieved in 5 days, and another 2-3 days may be necessary if plasmid curing was needed (Fig. 2*A* and *SI Appendix*, Fig. S3). The workflow included the design and synthesis of sgRNA and primers, followed by the construction and transformation of editor plasmid, the induction of *Sp*Cas9M-sfGFP, and the editing verification of *C. crescentus* clones. The *Sp*Cas9M fluorescigenic clones were identified before the colony PCR by a blue-light lamp (Fig. 2*A*). We have prepared a set of starting plasmids with different promoters and selection markers (Fig. 2*B*). Moreover, by replacing *sfgfp* with the chloramphenicol acetyltransferase coding gene (*cat*), an alternative CRISPR/*Sp*Cas9M-reporting plasmid (pSJX022) was constructed (Fig. 2*A* and *B*). Indicated by the chloramphenicol-resistance, clones expressing the full-length *Sp*Cas9M were identified and further confirmed by colony PCR and sequencing, showing comparable AEEs compared with the fluorescent reporting system (Fig. 2*C*).

To further streamline our CRISPR/*Sp*Cas9M editing system, other elements that may affect genome editing were characterized using the non-reporting plasmid pSJX005. This plasmid allowed us to exclude the dominant effects of the reporting proteins on the genome editing efficiency (Fig. 1*A*). We first evaluated the influence of varying lengths of H-arms on the editing efficiency in *C. crescentus*. Previous studies indicated that H-arm length could affect the HR ratio significantly in multiple bacteria (24). In this test, the *spmX* gene was chosen as the target gene and 0.5 µM vanillate (1/100 of the FUC in *C. crescentus*) was used to drive P*^van^*. With elongated H-arms, the *spmX*-knockout ratio increased gradually and reached 40.3% when the unilateral length of H-arms was 2,000 bp. No editing was detected when the length was less than or equal to 200 bp (Fig. 2*D*). These findings confirmed the weak HR activity in *C. crescentus* and demonstrate that H-arm length is a pivotal factor in regulating the editing efficiency. Nevertheless, the extensively long H-arms will lead to oversized plasmids, which might not be ideal for DNA assembly and transformation. Therefore, the unilateral length of H-arms utilized in subsequent study was 2,000 bp unless other indicated.

We also compared the editing efficiency for different promoters P*^van^* and P*^xyl^*, with no promoter as the control. The working concentration, 0.5 µM vanillate and 0.0003% (m/V) xylose, was used in this study. No significant difference of genome editing efficiency was observed between P*^van^* and P*^xyl^*, indicating a minimal discrepancy with the use of these two promoters (Fig. 2*E*). No edit was detected when the promoter was absent, confirming the genome editing is a Cas-dependent process (Fig. 2*E*).

Single guide RNA is another essential component in the CRISPR/*Sp*Cas9M system, which determines the cleavage specificity and also the editing efficiency. To investigate the influence of sgRNA on genome editing, five sgRNAs targeting different sites in *spmX* were designed by CHOPCHOP, a specialized web tool developed by Eivind Valen’s group (25). The results showed that editing efficiency varies from 20% to 40%, suggesting the sgRNA has a significant impact on the genome editing efficiency in our system (Fig. 2*F*).

Collectively, in addition to the CRISPR/*sp*Cas9M-reporting system, these results indicate that the use of a relatively long H-arm and the design of multiple sgRNAs are necessary for efficient genome editing in *C. crescentus*.

### Generation of gene deletion strains affecting asymmetric cell division

In *C. crescentus*, asymmetric cell division is modulated by a kinase and phosphatase pair, DivJ and PleC. The asymmetrical localization of DivJ/PleC at opposing cell poles regulates the cellular phosphorylation gradient and determines the fates of the daughter cells (26–28). Nevertheless, the polar accumulations of DivJ and PleC are not active processes but recruited by distinct scaffold proteins (Fig. 3*A*). Previous reports have shown that DivJ localization at the stalked cell pole was recruited by the PopZ-SpmX scaffold complex (29), while PleC localization at the swarmer cell pole was recruited by scaffold PodJ (30–32).

**Fig. 3.**
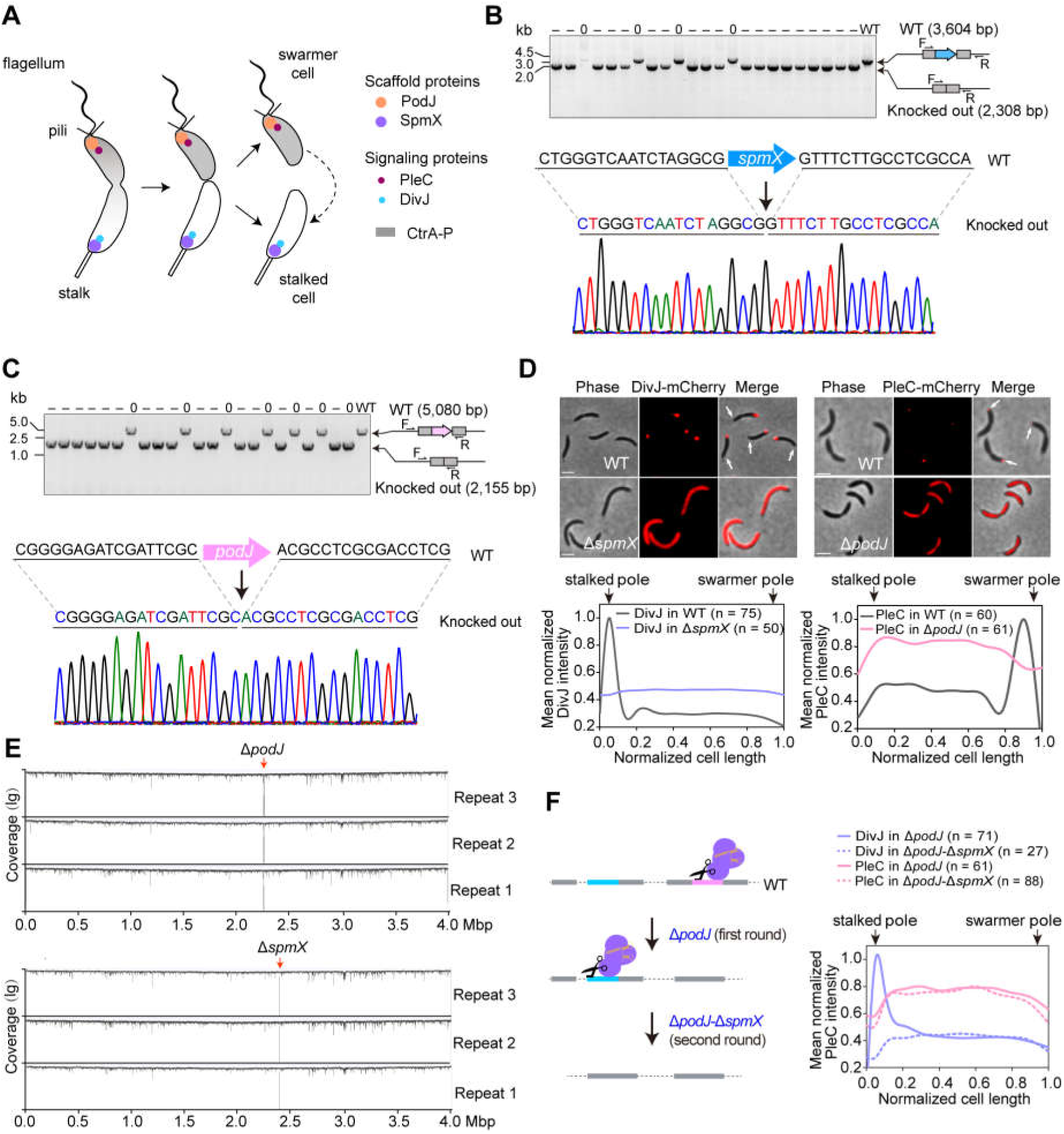
Generation of two scaffold gene knockouts that impairs the cell asymmetry. (*A*) Schematic of asymmetric cell division of *C. crescentus*. Two scaffold proteins, SpmX and PodJ, asymmetrically accumulate at the stalked cell pole and swarmer cell pole, recruiting signaling proteins DivJ and PleC, respectively. The localization of DivJ/PleC at the opposite cell poles regulates the downstream signal pathways and determines the cell fates. (*B*) Identification and confirmation of the *spmX*-knockouts by colony PCR and DNA sequencing. The *spmX*-knockouts were generated by CRISPR/*Sp*Cas9M-sfGFP system. (*C*) Identification and confirmation of the *podJ*-knockouts by cloning PCR and DNA sequencing. The *podJ*-knockouts were generated by CRISPR/*Sp*Cas9M-sfGFP system. (*D*) In-frame deletion of *spmX* and *podJ* disrupts the asymmetric localization of DivJ-mCherry and PleC-mCherry, respectively. Quantitative analysis of the cellular distributions of DivJ-mCherry and PleC-mCherry in Δ*spmX* or Δ*podJ* mutants, using the wild-type (WT) cells as the controls. Data were normalized with the highest intensity as 100% in each test combination. The white arrows indicate the swarmer cell poles. All scale bars, 2 μm. (*E*) Whole genome sequencing suggests no off-targets in Δ*spmX* or Δ*podJ* mutants. The sequencing reads obtained from each sample were mapped to the reference genome of *C. crescentus* NA1000. The grey lines indicate the unmapped ratio for each 100 bp calculated window in the chromosome. The localization sites of *podJ* and *spmX* are indicated by red arrows. Three biological repeats for each knockout were tested. (*F*) Cell asymmetry was completely lost after mutation of both *spmX* and *podJ.* A flow chart of double mutation of *spmX* and *podJ* using the CRISPR/*Sp*Cas9M-reporting system (left panel). Quantitative analysis of the cellular distributions of DivJ-mCherry and PleC-mCherry in Δ*podJ*-Δ*spmX*, using the first round of Δ*podJ* cells as the controls. Data were normalized with the highest intensity of DivJ-mCherry in Δ*podJ* as 100%.

To genetically confirm these interactions, the coding genes for SpmX and PodJ were in-framely knocked out and systematically analyzed using the CRISPR/*Sp*Cas9M-reporting system in *C. crescentus.* In total, 19 out of 23 of clones were successfully edited in the *spmX* deletion experiment, with AEE values up to 82.6% (Fig. 3*B*). Meanwhile, an AEE of 73.9% was obtained in the *podJ* deletion experiments, supporting an efficient and practical editing method for our system (Fig. 3*C*). The DivJ and PleC proteins were labeled by mCherry and transformed in these markless deletion cells, respectively. Their localizations were then analyzed using inverted fluorescence microscopy. Indicated by the polar stalk, we observed that DivJ-mCherry specifically accumulated at the stalked cell poles while PleC-mCherry specifically accumulated at the swarmer cell poles in the wild-type strains. Nevertheless, DivJ-mCherry was unable to localize at the cell poles when *spmX* was deleted. Similar disperse of PleC-mCherry in the cells was observed when *podJ* was deleted (Fig. 3*D*). These observations are consistent with the previous reports, suggesting these scaffold proteins are necessary for DivJ/PleC asymmetric localization.

Possible off-target editing was analyzed by whole genome sequencing after curing of the editor plasmid. Three clones each from Δ*spmX* and Δ*podJ* were randomly selected for genome sequencing, using an Illumina HiSeq TM 2000 platform. Approximately 1 gigabyte data was obtained for each clone, providing ∼200× average sequencing depth and 100% genome coverage. No off-targets were observed for any of the six clones, indicating no viable nonspecific cleavage in our system (Fig. 3*E*).

To evaluate the robustness of our CRISPR/*Sp*Cas9M-reporting system, a double mutation strain was constructed by sequential knocking-out of *podJ* and *spmX*. The first round of *podJ* deletion was completed in 5 days, followed by plasmid curing for 3 days. The *spmX* deletion editor plasmid was then electroporated and the genome editing was achieved in another 5 days. The localization of DivJ-mCherry and PleC-mCherry in the double mutant Δ*podJ*-Δ*spmX* showed a complete loss of localization asymmetry of the two signaling proteins (Fig. 3*F* and *SI Appendix*, Fig. S4*A*). Compared with that in single mutation, the AEE of the second-round editing in double mutation did not change significantly (*SI Appendix*, Fig. S4*B*), demonstrating that our system that could be used for multi-site genome editing.

We also conducted a set of gene knockouts to assess the performance consistency of the CRISPR/*sp*Cas9M-reporting system. Nine genes of varying sizes (0.9-7.7 kb) were selected as targets by designing three different sgRNAs for each. The results demonstrated that all nine genes could be readily knocked out, though variable AAEs were obtained (*SI Appendix*, Fig. S4*C*). It’s worth noting that *zitP* and *pleC* among these genes are hardly knocked out if we use the non-reporting system (data not shown). Hence, these results suggest our CRISPR/*Sp*Cas9M-reporting system has a reliable performance in editing different genome loci.

### Achieve targeted genome insertion in *C. crescentus*

Scientists have been unable to achieve in-frame and markless gene insertion in *C. crescentus* to date. In order to solve this problem, the CRISPR/*Sp*Cas9M-reporting system was employed to test the targeted genome insertion in *C. crescentus*.

We first designed an in-frame insertion by replacing *spmX* with the mCherry encoding gene. A pair of H-arms with *mcherry* in the middle were assembled into the pSJX016 vector, and two different sgRNAs were designed to avoid potential low AEE (Fig. 4*A*). The editor plasmid was then transferred into *C. crescentus* through electroporation. After exclusion of the *sp*Cas9M mutants, the remaining clones were verified by colony PCR and further confirmed by sequencing (Fig. 4*B*). Quantitative analysis showed that the AEEs of the two sgRNAs were 80.6% and 52.8%, respectively, demonstrating this approach is an efficient insertion method (Fig. 4*C*).

**Fig. 4.**
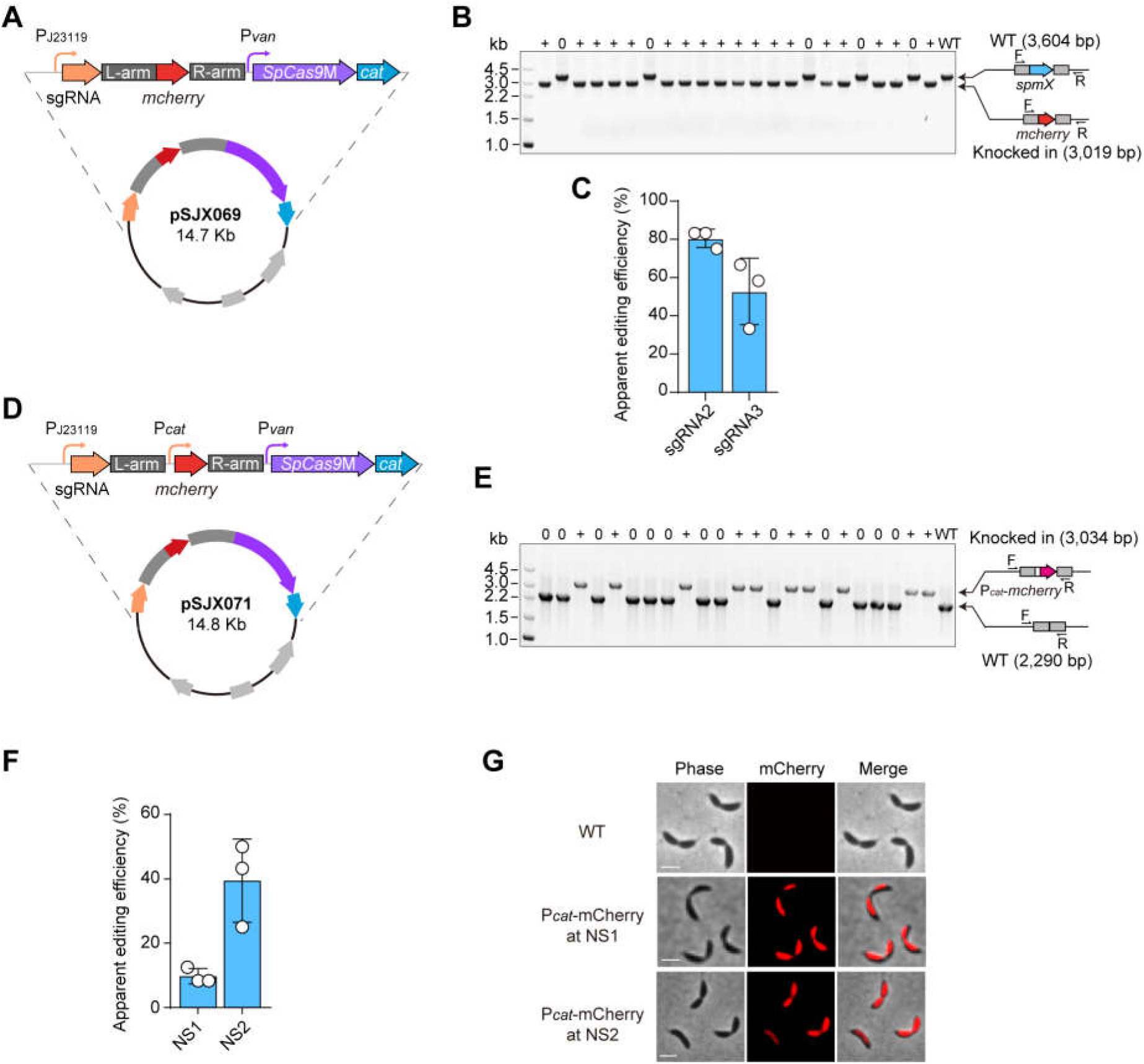
Targeted genome insertion in *C. crescentus* by CRISPR/*Sp*Cas9M-reporting system. (*A*) A schematic diagram of the plasmid used in targeted genome replacement. The mCherry coding sequence was designed in the middle of H-arms, aiming to make an in-frame replacement of *spmX* in the chromosome. The chloramphenicol acetyltransferase coding gene, *cat*, was used as the reporter gene. (*B*) Identification of the *mcherry* replacements by colony PCR. (*C*) Quantitative analysis of the apparent editing efficiency of *mcherry* replacements mediated by two different sgRNAs targeting *spmX*. The data are presented as means ± SEM from 3 independent biological replicates. A total of 36 clones (*n*) were tested for each sample. (*D*) A schematic diagram of the plasmid used in targeted genome insertion. The P*^cat^* promoter and the mCherry coding sequence were designed in the middle of H-arms, which were targeted to insert into the neutral insertion sites in *C. crescentus* genome. (*E*) Identification of the P*^cat^* -*mcherry* insertions by colony PCR. (*F*) Quantitative analysis of the apparent editing efficiency of P*^cat^*-*mcherry* insertions. NIS1 and NIS2 are the two different neutral insertion sites tested in this study. The data are presented as means ± SEM from 3 independent biological replicates. A total of 36 clones (*n*) were tested for each sample. (*G*) Phenotypic analysis confirms P*^cat^*-*mcherry* insertion and indicates that the fitness of *C. crescentus* was not affected by the insertions. All scale bars, 2 μm.

When using this method for applications such as gain of function or genome barcoding, a <u>n</u>eutral insertion <u>s</u>ite (NIS) may be necessary to avoid polar effects (33). In this study, we have identified a set of putative NISs in *C. crescentus* by using the targetFinder algorithm (*SI Appendix*, Table S2). The top two NISs (NIS1 and NIS2) were selected as targets to evaluate the corresponding AEEs in this study. A construct containing a sgRNA and a P*cat*-*mcherry* sequence arranged between two H-arms was built based on pSJX016 vector (Fig. 4*D*). The editor plasmid was then transferred into *C. crescentus* and the clones were selected and detected by colony PCR. A relatively higher AEE was observed for NIS2 when compared to NIS1, reaching up to 40%. This result indicates NIS2 could be an appropriate spot for gene insertion in *C. crescentus* chromosome (Fig. 4*E* and *F*). The red fluorescence of mCherry was observed using inverted fluorescence microscopy, which confirmed the correct insertion and expression of *mcherry* (Fig. 4*G*). Moreover, no detectable morphological changes were observed for all the edited cells (Fig. 4*G*), indicating that the insertion spots resulted in no obvious disruption to the fitness of *C. crescentus*.

### Applying the CRISPR/*Sp*Cas9M-reporting system to other α-proteobacteria

As close relatives of *C. crescentus*, *S. meliloti* and *A. fabrum* are plant symbiont and pathogen, respectively, and both play critical roles in modern agricultural biotechnology (34). *S. meliloti* is also used as a cell factory to produce native compounds recently (22, 35). Nevertheless, both *S. meliloti* and *A. fabrum* are hard-to-edited organisms due to the possible lack of cellular HR activity. Genome editing tools such as CRISPR-mediated base editing and transposon have been developed for *A. fabrum* and *S. meliloti*, respectively (21, 22, 36). However, in-frame gene deletion or insertion remains a significant challenge.

As a proof-of-concept application, we investigated the editing potential of the CRISPR/*Sp*Cas9M-reporting system in *A. fabrum* and *S. meliloti*. The *tdk* gene encoding thymidine kinase was selected as the target in this study. Thymidine kinase confers the cell sensitivity to 5-fluoro-2′-deoxyuridine (5-FudR) by phosphorylating 5-FudR to the toxic thymidine analog fluoro-dUMP (F-dUMP) (37–39). In *A. fabrum*, the corresponding H-arms and sgRNA were cloned to the pSJX016 vector as used in *C. crescentus.* Three different sgRNAs targeting *tdk* were designed and their knockout efficiencies were evaluated (Fig. 5*A* and *B*). Colony PCR and sequencing results showed that the AEEs of the three sgRNAs were 86.7% (sgRNA1), 29.9% (sgRNA2), and 8.3% (sgRNA3). To confirm the phenotype of these edited cells, a growth experiment was performed on peptone yeast extract (PYE) plates containing 200 μg/ml 5-FudR. As expected, the Δ*tdk* cells survived on the PYE plates in the presence of 5-FudR, while the wild-type *A. fabrum* cells could not (Fig. 5*C*).

**Fig. 5.**
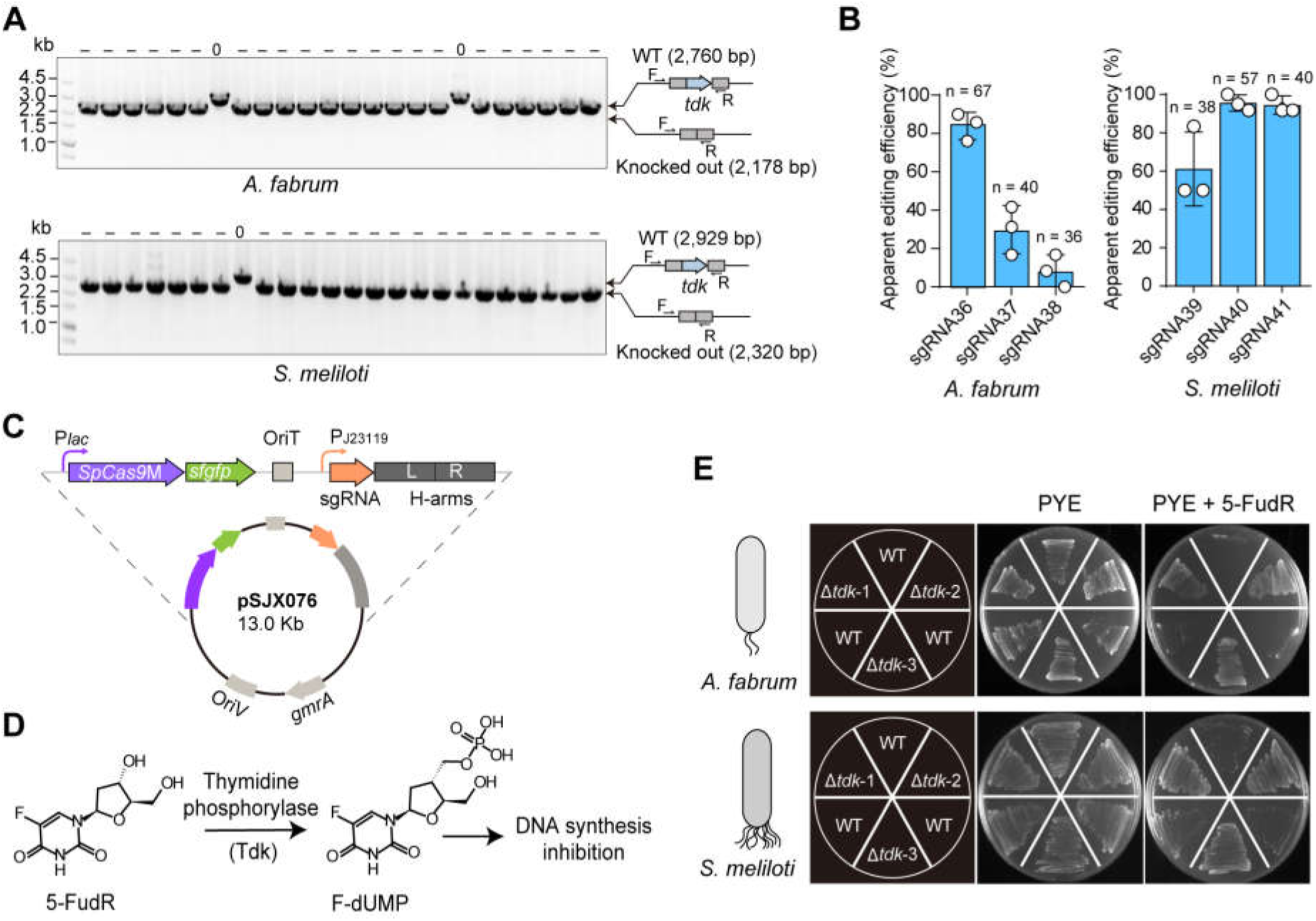
Apply the CRISPR/*Sp*Cas9M-reporting system to *A. fabrum* and *S. meliloti*. (*A*) Identification of the *tdk*-knockouts in *A. fabrum* and *S. meliloti* by colony PCR. (*B*) Quantitative analysis of the apparent editing efficiency of the *tdk* deletion. Three sgRNAs targeting *tdk* were designed and tested for each strain. The data are presented as means ± SEM from 3 independent biological replicates. (*C*) Schematic diagram of the conjugation plasmid used for genome editing in *S. meliloti*. The inducible promoter P*^lac^* was used to drive *sp*Cas9M expression. (D) Phenotypic analysis confirms the deletion of *tdk.* The *tdk* encoded thymidine kinase functions in catalyzing 5-FudR to the toxic F-dUMP, which inhibits DNA synthesis. The correct Δ*tdk* mutants could survive on the PYE plates in the presence of 200 μg/ml 5-FudR. Representative plates from three independent experiments are shown after being cultured at 30 °C for 3 days.

A similar experimental design was used for *S. meliloti*, but in this case the conjugation transformation method was used. The CRISPR/*Sp*Cas9M-reporting system, which included a P*^lac^*-driven *Sp*Cas9M, was cloned into a pK18mob plasmid containing an *oriT* DNA region. The final editor plasmid pSJX072 can be feasibly transformed into *S. meliloti* (Fig. 5*C*). The clones selected from PYE plates were screened by colony PCR followed by sequencing. Relatively high AEEs were shown for all the three sgRNAs, reaching 61.1%, 95.6%, and 94.5%, respectively (Fig. 5*A* and *B*). The growth experiment confirmed only the Δ*tdk* edited cells could survive on the PYE plates in the presence of 5-FudR (Fig. 5*D*).

Together, these results suggest that the CRISPR/*Sp*Cas9M-reporting system is practical and reliable in both *A. fabrum* and *S. meliloti*, demonstrating this system as an efficient, in-frame, and markless editing tool.

## Discussion

The CRISPR/Cas system has become the dominant genome editing method in a wide variety of organisms by virtue of its efficiency, conveniency, and reliability. However, its success depends on either HR or non-homologous end joining (NHEJ) activity in cells since repair of chromosomal DSBs caused by Cas cleavage are necessary. In bacteria, where NHEJ pathways are typically absent, most CRISPR/Cas genome engineering tools rely on HR-based repair. In the current study, we developed a HR-assisted CRISPR/Cas9 system by enhancing the HR activity in *C. crescentus*. Moreover, the genome editing efficiency was significantly improved by systematic optimization of the Cas9 codon usage and expression level, and uniquely, by introducing a reporting system. The resulting CRISPR/*Sp*Cas9M-reporting system allows for efficient, markless, and rapid genome editing in *C. crescentus*, offering multiple advantages over existing tools (Table 1). We successfully applied this system to two agricultural bacteria, *S. meliloti* and *A. fabrum*, and the results of these experiments suggest our CRISPR/*Sp*Cas9M-reporting system could serve as a general editing strategy for other challenging organisms.

**Table 1.** Comparison of CRISPR/*Sp*Cas9M-reporting system with other genome editing tools in *C. crescentus*.

| Genome editing method | Mutation type | Editing efficiency | Selection marker | Time cost | Reference |
| --- | --- | --- | --- | --- | --- |
| Counterselection | Knock out, knock in | Very low | Markless | >3 weeks | Ely, et, al. 1991(17) |
| Genome integration | Knock in | 90-100% | With markers | ~1 week | Martin, et, al. 2007(19) |
| CRISPRi | Knock down | 90-100% | With markers | ~1 week | Mathilde Guzzo, et, al. 2020(20) |
| CRISPR/*SpCas9M* reporting system | Knock out, knock in | 20-90% | Markless | ~1 week | This study |

In this study, the most important factor in achieving reliable edits at distinct genome loci or in differnt organisms was the development of a reporting system after discovering the CRISPR escapers. In fact, CRISPR escape is a widespread phenomenon that has been reported in multiple organisms including bacteria (40, 41). For example, studies in *Corynebacterium glutamicum* showed that escapers arose from the recombination of repeat sequences between the CRISPR arrays, which deleted the spacer region on the sgRNA(42). In *Escherichia coli*, studies reported that the escapers might be caused by the mutation of targeted DNA. These studies hypothesized that the SOS response system is activated by RecA after Cas9 cleavage, which turns on the expression of error-prone DNA polymerase and leads to mutations in the target DNA (43, 44). However, the escapers were still observed after knocking out *recA* and the DNA polymerase genes, suggesting that escape mechanisms in bacteria remain complex (44, 45). Recently, Li et al. identified frequent insertions of insertion sequence 5 (IS5) in *cas9* as the primary cause of escapers in *E. Coli*, while the triggering factor was elusive (41). Our findings in *C. crescentus* support a novel type of CRISPR escapers generated by random deletions within the *Sp*Cas9 coding sequence. The mutated plasmids were survived from the potential anti-CRISPR defenses, indicating that some unrecognized repair systems may be involved in this process. Although the exact mechanisms underlying CRISPR escaper are not fully understood, we developed a practical CRISPR/Cas-reporting system based on these observations. By leveraging reporter genes, we enabled visual detection of *Sp*Cas9M expression in *C. crescentus*, which greatly enhanced the apparent genome editing efficiency. Importantly, this system performed consistently across various gene targets and facilitated in-frame insertion and iterative editing, as well.

Despite these improvements, the AEE did not reach 100% even with the reporting system in place. Some clones exhibited fluorescent signals or antibiotic resistance but the intended edits were missing. This outcome may be explained by two possibilities. One possibility is that *Sp*Cas9M undergoes frameshift mutations, rendering it inactive for cleavage. We have observed some fluorescent clones that contained frameshift-mutated *Sp*Cas9M by sequencing (data not shown). The other possibility is that the sgRNA might not efficiently guide *Sp*Cas9M to the cleavage site. This finding underscores the importance of using professional design software and testing multiple sgRNAs to optimize genome editing success.

## Materials and methods

### Bacterial strains and culture conditions

Detailed information on the growth conditions of *E. coli*, *C. crescentus* NA1000, *A. fabrum* C58, and *S. meliloti* 1021 are provided in *SI Appendix, Materials and Methods*. Bacterial strains used in this study are detailed in *SI Appendix,* Table S3.

### Phylogenetic analysis

All the 16S rRNA sequences were downloaded from the NCBI database. The sequences of 16S rRNA were aligned by ClustalW (46) and the phylogenetic tree was built by MEGA7 (47). The details are provided in *SI Appendix, Materials and Methods*.

### Editor plasmid construction

Details on the construction of editor plasmids are provided in *SI Appendix, Materials and Methods*. The lists of plasmids, sgRNAs, and oligonucleotides are provided in *SI Appendix,* Tables S4-6.

### Bacteria transformation

The transformation of *E coli* used a chemical method as described by (48). The transformation of *C. crescentus* and *A. fabrum* used an electroporation method (17) and the transformation of *S. meliloti* used a conjugation method (49). The competent cells were prepared as described in *SI Appendix, Materials and Methods*.

### Genome editing analysis

PCR and Sanger sequencing were used for genome editing analysis. The details of genome editing analysis and plasmid curing are provided in *SI Appendix, Materials and Methods*.

### Off-target analysis by whole genome sequencing

Whole genome sequencing was performed on these samples using the Illumina HiSeq TM 2000 platform. The raw data were processed by Fastp (version 0.19.7) (50) and Geneious Prime software (version 2019). Details on analysis are given in *SI Appendix, Materials and Methods*.

### NIS analysis

The putative neutral insertion sites were predicted by targetFinder algorithm (51). Details of NIS analysis are given in *SI Appendix, Materials and Methods*.

### Inverted fluorescence microscopy

Protein localization analysis was performed as previously described (52, 53). Details are provided in *SI Appendix, Materials and Methods*.

### Data availability

Data supporting the findings of this study are available within the main Manuscript and the *SI Appendix, Materials and Methods*. The whole genome sequencing data generated in this study were uploaded to the SRA database (PRJNA1187698).

## Supporting information

Supporting Information for A CRISPR/SpCas9M-reporting system for efficient and rapid genome editing in Caulobacter crescentus

## ACKNOWLEDGMENTS

This work is supported by the National Key R&D Program of China (grant no. 2024YFA1700128 to W. Z.), the Strategic Priority Research Program of the Chinese Academy of Sciences (grant no. XDB0480000 to W. Z.), the National Natural Science Foundation of China (grant no. 32471494 and U22A201247 to W. Z.), and the Guangdong Basic and Applied Basic Research Foundation (grant no. 2023A1515030069 to W. Z.).

## Notes

### Competing Interest Statement

The authors have declared no competing interest.

