## Supporting Information for A CRISPR/SpCas9M-reporting system for efficient and rapid genome editing in Caulobacter crescentus for "A CRISPR/*Sp*Cas9M-reporting system for efficient and rapid genome editing in *Caulobacter crescentus*"

**This PDF file includes:**

Supplementary Material and Methods

Figures S1 to S4

Tables S1 to S6

SI References

### Supplementary Material and Methods

#### Bacterial strains and culture conditions.

All strains, plasmids, sgRNAs, and oligonucleotides used in this study are listed in *SI Appendix,* TableS 3-6. Three α-proteobacteria, i.e., *C. crescentus* NA1000, *A. fabrum* C58 and *S. meliloti* 1021, were used as the wild-type strains in this study. All α-proteobacteria and their derivatives were grown in PYE liquid medium or on PYE agar plate at 30 °C. The *E. coli* DH5α cells were cultivated in Luria-Bertani (LB) medium at 37 °C.

When necessary, media were supplemented with antibiotics at the following concentrations (liquid/solid media for *C. crescentus*; liquid/solid media for *A. fabrum*; liquid/solid media for *S. meliloti*; liquid/solid media for *E. coli*, in µg/mL): kanamycin (5/25; 50/50; 50/50; 50/50), spectinomycin (25/50; 50/100; 600/600; 50/50), chloramphenicol (1/2; 30/30; 30/30;30/30), and gentamicin (1/5; 20/20; 50/50;50/50). The electroporation of *C. crescentus* and *A. fabrum* followed the method described in (1), while the conjugation of *S. meliloti* followed the method described in (2). The transformation of *E coli* used a chemical method as described by (3).

Two different inducible promoters, $P_{xyl}$ and $P_{van}$, were used to manipulate the expression level of *Sp*Cas9M. Xylose and vanillate are the inducers of $P_{xyl}$ and $P_{van}$, respectively. The frequently-used concentration (FUC) of xylose is 0.03% (m/V), and the FUC of vanillate is 50 µM.

#### Phylogenetic analysis

To analyze the evolutionary relationship between *C. crescentus* and its relatives within α-proteobacteria, the DNA sequences of 16S rRNA from α-proteobacteria, including *C. crescentus*, *A. fabrum*, and *S. meliloti* were downloaded from the NCBI database, and aligned by ClustalW (4). A phylogenetic tree was built by MEGA7 (5) using 37 representative 16S rRNA sequences with the Neighbor-Joining method (bootstrap: 1,000).

#### Editor plasmid construction

All editor plasmids were constructed using the Gibson assembly method (6) based on the pBXMCS-2 vector. The Cas genes with condon optimization including *Sp*Cas9M, *Fn*Cas12aM, *Sth1*Cas9M, and *Sth3*Cas9M were synthesized by Sangon Biotech (Shanghai, China). The sgRNAs used in this study were designed using CHOPCHOP (7) and synthesized by Sangon Biotech. The corresponding H-arms were cloned from bacterial total DNA. When necessary, the reporter gene *sfgfp* or *cat* were fused to the C-terminus of *Sp*Cas9M, using a flexible linker "HRSAT". All DNA fragments including the promoters were cloned into pBXMCS2 or pBVMCS6 vector, and were named as all-in-one plasmids. We prepared a set of starting editor plasmids with different selection markers and promoters (Fig. 2*B*). When targeting a new site, these plasmids could be used as a start point by replacing the sgRNA and H-arms in just one Gibson assembly.

**Preparation of competent cells and electroporation of *C. crescentus* and *A. fabrum***

To prepare the competent cells of *C. crescentus* and *A. fabrum* for electroporation, the corresponding bacterium from glycerol stocks was inoculated on PYE plates at 30 °C for around 2-3 days. A mono-clone was selected and inoculated in 3 mL PYE liquid medium and grown overnight at 30 °C, 200 rpm. Then, a 1 mL culture was transferred into 100 mL PYE liquid medium and cultured to a final $OD_{600}$ around 0.7-1.0 at 30 °C. The culture was collected in a sterilized ice-cold 50 mL centrifuge tube and centrifuged at 6,000 *g* for 10 min, 4 °C. The supernatant was discarded and the pellets were resuspended in 40 mL cold distilled water, followed by centrifugation at 6,000 *g* for 10 min, 4 °C. Pellets were washed three times and were then resuspended in 1 mL cold distilled water and further separated into 100 μL aliquots.

For electroporation, 1 μg editor plasmid was first mixed with 100 μL competent cells for 20 min on ice. Then, the mixture was transferred to an ice-cold Bio-Rad Gene Pulser cuvette (Richmond, CA) and subjected to a 2.5 kV electric shock. The capacitor was set at 25 μF and the resistance was set at 200 ohms. After the electric shock, cells in the cuvette were transferred immediately to a 1.5 mL tube containing 1 mL of PYE. After a 3-hours recovery period at 30 °C, the *C. crescentus* cells were plated on the appropriate selective PYE plates containing a corresponding inducer. For *A. fabrum*, all electroporation procedures were the same as were used for *C. crescentus*, except using 1-hour recovery time.

#### The conjugation of *S. meliloti*

The editor plasmid was first transformed into the donor strain WM6026. WM6026 is an auxotrophic *E. coli* strain whose growth relies on exogenously supplemented Diaminopimelic acid (DAP). The overnight WM6026 culture was diluted 100-fold in fresh LB medium containing gentamicin and DAP (100 μg/mL). A similar dilution was performed for the recipient of *S. meliloti*, using LB medium without antibiotics. The donor and recipient cells were grown to the exponential phase ($OD_{600}$ = 0.4-0.6) and mixed by volume at a 1:3 ratio. After centrifuging at 6,000 *g* for 5 min, the pellets were then resuspended in 150 μL of LB liquid medium and spotted on LB plates with DAP. The cells were incubated at 30 °C overnight, recovered with a sterile spatula, and resuspended in 1 mL LB liquid medium at 30 °C for 2 h. Finally, these transconjugants were selected by gentamicin on LB plates containing a corresponding inducer.

#### Genome editing analysis

The blue-light lamp (DUT-48, DC24V, 0.8A) or the PYE plates with chloramphenicol (2 μg/mL) were use to identify the clones normally expressing *Sp*Cas9M. The clones exhibiting green fluorescence under the blue-light lamp or growing on the PYE plates with chloramphenicol, indicating the *Sp*Cas9M was normally expressed in cells, were the potential editing strains.

The colony PCR was used to evaluate the genome editing efficiency or apparent editing efficiency. The single clone was picked from plates and suspended in 20 μL reaction mixtures contained 10 μL of Green Taq Mix (Vazyme Biotech), 1 μM primer and 8 μL of $H_2O$. Primer pairs were designed at upstream of the left homologous arm and downstream of the right homologous arm. PCR products were separated by 1% agarose gels and stained with SYBR Safe (Thermo Scientific). Negative controls (usually the wild-type cells) and experimental samples were analyzed in parallel to identify non-specific bands. The PCR products were sent to Sangon Biotech for sequencing analysis.

**Editor plasmid curing**

The correct editing clone of *C. crescentus* was cultured in 5 mL PYE liquid medium at 30 °C, without any antibiotics. After 15-18 hours, the cells were collected and diluted 1:1000 in fresh PYE medium. About 50 µL of the diluted culture was spread on PYE plates without antibiotics, and then incubated at 30 °C for about 2 days. Single clones were picked and replica-spotted on PYE plates with no antibiotic, as well as on plates with antibiotics. The clones that grew only on the antibiotic-free plates were selected as the cured strains.

**Off-target analysis by whole genome sequencing**

To determine whether the off-targets occurred in the edited strains, the verified clones were collected after curing the editor plasmid. Whole genome sequencing was performed on these samples using the Illumina HiSeq TM 2000 platform. The raw data were pre-processed and the low-quality data were filtered using Fastp (version 0.19.7) (8). The final data obtained from each edited strain was mapped to the reference genome sequence of *C. crescentus* NA1000 and the genome variations between edited strain and NA1000 were analyzed by Geneious Prime software (version 2019). The possible variations were further verified by Sanger sequencing.

**5-FudR assay**

To confirm the phenotype of *tdk* deletions in *A. fabrum* and *S. meliloti*, the *tdk*-knockouts and the corresponding wild-type cells were cultured on PYE plates containing 200 µg/ml 5-FudR. The PYE plates without 5-FudR were used as controls. These cells were cultured on plates at 30 °C for 3 days. Three repeats were performed for each assay.

**NIS analysis**

The NIS analysis was performed for *C. crescentus* genome by using the targetFinder algorithm (https://github.com/ECBCgit/targetFinder). The genome sequence of *C. crescentus* was analyzed, and the convergently transcribed genes within a range of gap sizes (300 to 2000 bp used here) were identified. The gaps between the interval of convergently transcribed genes were then further screened by additional annotations, including local repetitive structures, larger repetitive structures, and additional repetitions (9). Finally, the remaining gaps were defined as putative phenotypically neutral insertion sites.

**Inverted fluorescence microscopy and protein localization analysis**

The imaging of fluorescent proteins in *C. crescentus* was performed as previously described (10). Cells were grown to the exponential phase in PYE medium, and then induced by the corresponding inducers at indicated concentrations for 2-3 hours. The cells were immobilized on a 1.5% agarose pad containing PYE medium and imaged with a Nikon Eclipse T*i*2-E inverted fluorescence microscope. Fluorescence of mCherry was detected using the TRITC filter (Nikon, excitation filter 542/20, dichroic mirror 570, and emission filter 620/52) or the Texas Red filter (Nikon, excitation filter 555/35, dichroic mirror 585, and emission filter 630/70). The fluorescence intensity along the cell length was quantitatively analyzed using MicrobeJ (11).

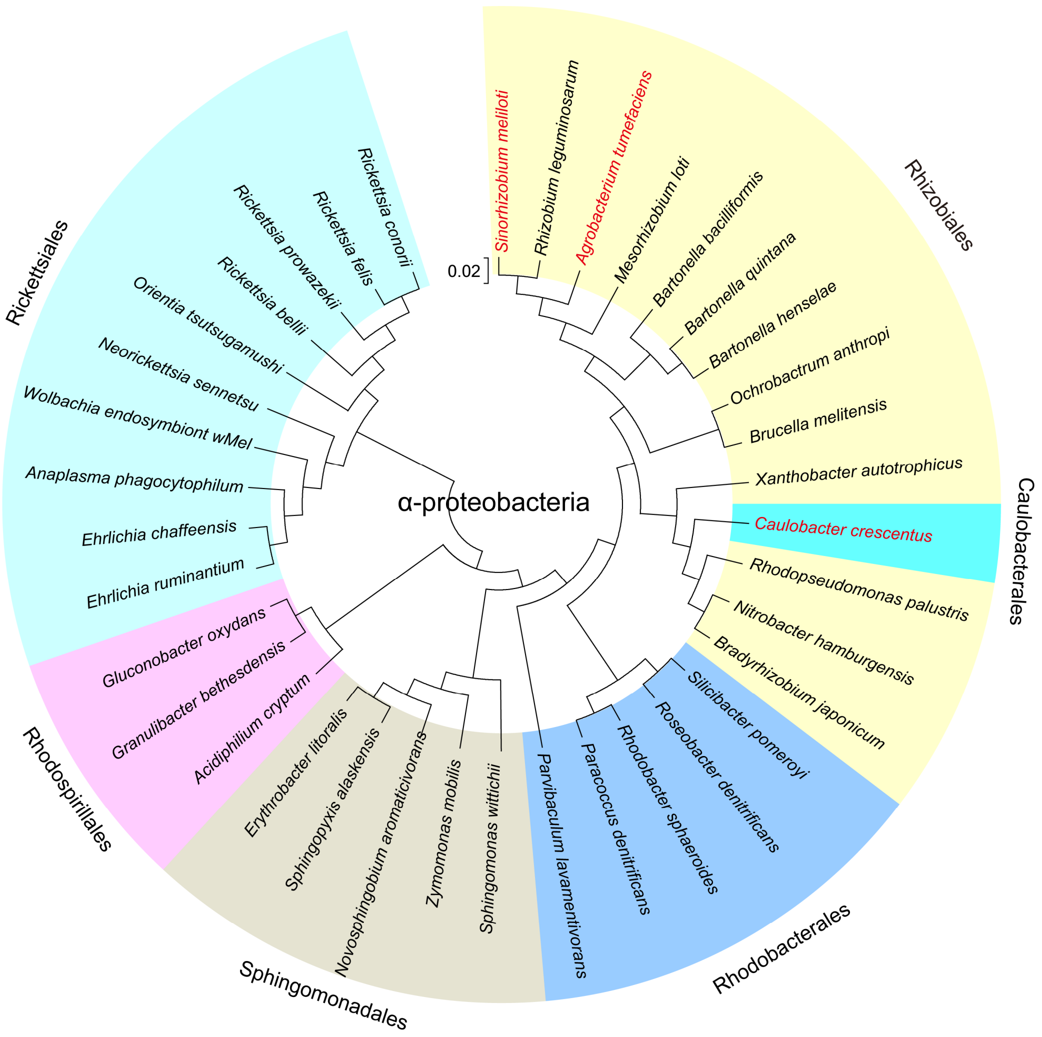


**Fig. S1. Phylogenetic analysis of *C. crescentus* in α-proteobacteria.**

Phylogenetic tree was built using the 16S rRNA sequences of representative α-proteobacteria by MEGA7 with Neighbor-Joining method (bootstrap: 1,000). A total of 37 α-proteobacteria were selected and the colors overlaid on these α-proteobacteria represent different orders. The NCBI accession numbers of 16S rRNA sequences used in this study are listed as below: NR_118988.1 (*Sinorhizobium meliloti*), NR_118339.1 (*Rhizobium leguminosarum*), NR_174322.1 (*Agrobacterium tumefaciens*), NR_025837.1 (*Mesorhizobium loti*), NR_119296.1 (*Bartonella bacilliformis*), NR_118693.2 (*Bartonella quintana*), NR_074335.2 (*Bartonella henselae*), HG917892.1 (*Ochrobactrum anthropi*), NR_043003.1 (*Brucella melitensis*), NR_074255.1 (*Xanthobacter autotrophicus*), M83799.1 (*Caulobacter crescentus*), NR_115542.1 (*Rhodopseudomonas palustris*), NR_074313.1 (*Nitrobacter hamburgensis*), NR_036865.1 (*Bradyrhizobium japonicum*), AF434674.2 (*Silicibacter pomeroyi*), NR_118769.1 (*Roseobacter denitrificans*),

KF791043.1 (*Rhodobacter sphaeroides*), NR_119264.1 (*Paracoccus denitrificans*), NR_029105.1 (*Parvibaculum lavamentivorans*), LC508800.1 (*Sphingomonas wittichii*), NR_028792.1 (*Zymomonas mobilis*), NR_118791.1 (*Novosphingobium aromaticivorans*), NR_115203.1 (*Sphingopyxis alaskensis*), OQ410310.1 (*Erythrobacter litoralis*), NR_119294.1 (*Acidiphilium cryptum*), NR_074276.1 (*Granulibacter bethesdensis*), NR_112534.1 (*Gluconobacter oxydans*), NR_074513.2 (*Ehrlichia ruminantium*), NR_074500.2 (*Ehrlichia chaffeensis*), NR_044762.1 (*Anaplasma phagocytophilum*), MK940242.1 (*Wolbachia endosymbiont wMel*), NR_074386.1 (*Neorickettsia sennetsu*), NR_025860.1 (*Orientia tsutsugamushi*), NR_074484.2 (*Rickettsia bellii*), NR_044656.2 (*Rickettsia prowazekii*), OM912382.1 (*Rickettsia felis*), and NR_074469.2 (*Rickettsia conorii*).

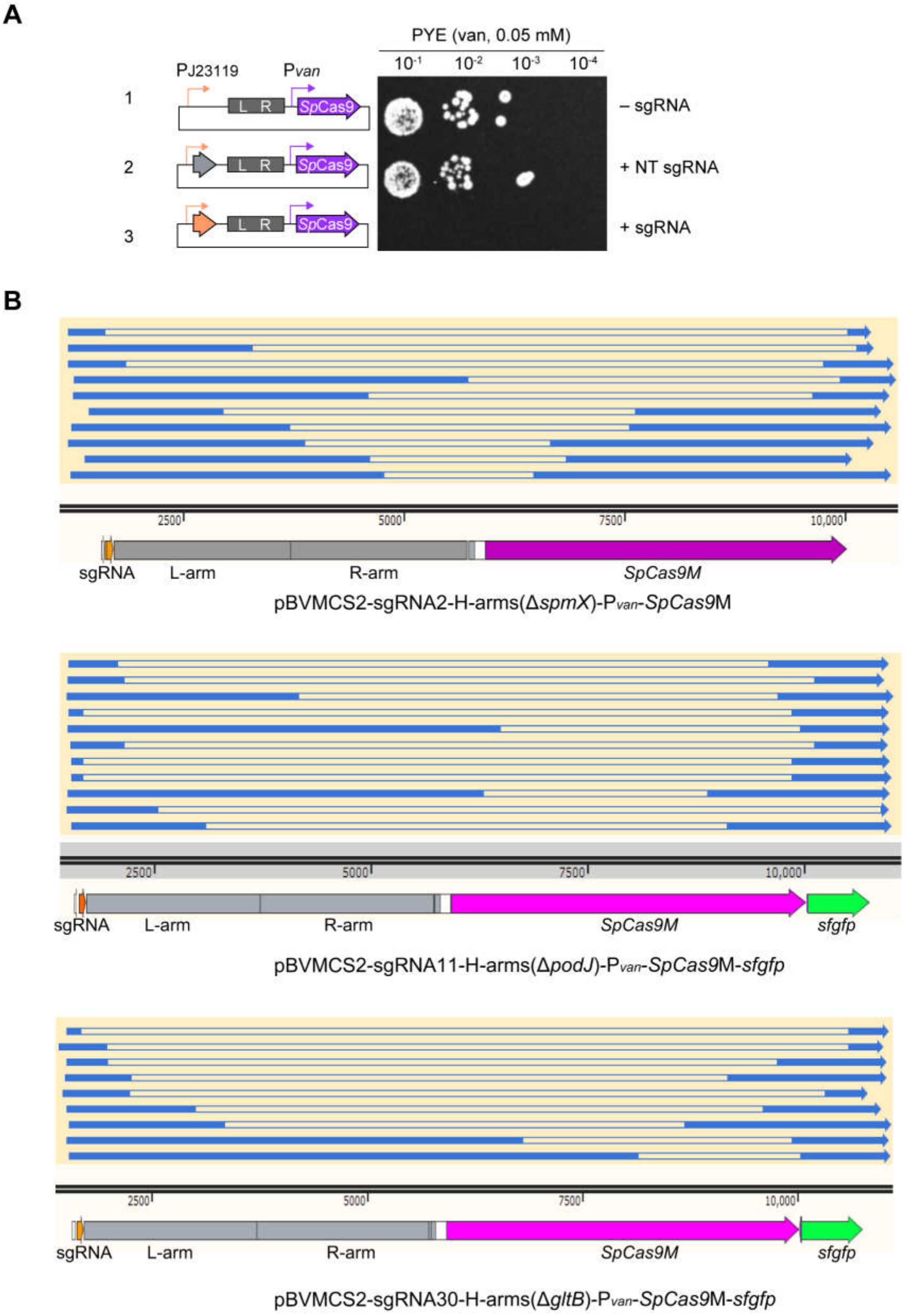


**Fig. S2. Analysis of the components that may affect the genome editing efficiency in HR-assisted CRISPR/Cas system.**

(*A*) The CRISPR/*Sp*Cas9 cleavage is lethal for *C. crescentus.* The plasmids containing

*Sp*Cas9 with or without sgRNA (left panel) were electroporated into the aliquots of *C. crescentus* competent cells, respectively. After a 3-hours recovery period, five microliters of serial dilutions were spotted onto the PYE plates that were supplemented with 50 μM vanillate. These plates were then observed and photographed after a 72-hours cultivation at 30°C (right panel). One representative of three plates was shown. The plasmid with non-targeting sgRNA (NT sgRNA) was used as the control. (*B*) The sequence analysis of editor plasmids in CRISPR escapers reveals that significant deletions on all the coding sequence of *Sp*Cas9M. The surviving colonies without editions were randomly selected from *spmX*, *podJ* and *gltB* editing experiments and their editor plasmids were sequenced by Sangon Biotech (Shanghai). These sequences were then aligned to the sequences of the starting editor plasmids, respectively. The lines filled no colors indicate the DNA sequences are absence in the editor plasmids.

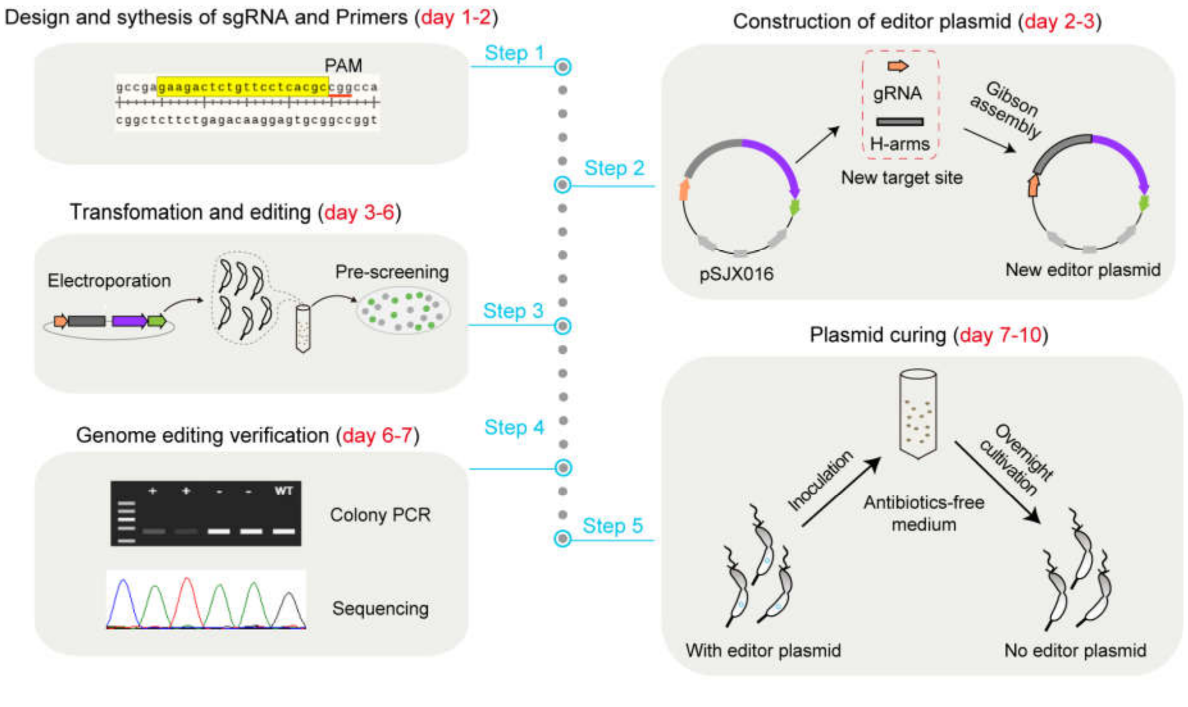


Genome editing workflow of CRISPR/*Sp*Cas9M-reporting system

**Fig. S3. The genome editing workflow of CRISPR/*Sp*Cas9M-reporting system in *C. crescentus*.**

First, the sgRNA and the corresponding primers are designed and synthesized (step one). Then, the new editor plasmid is constructed by replacing the old sgRNA and H-arms with the new sgRNA and H-arms in one Gibson assembly. The corrected editor plasmid could be checked by PCR and further by sequencing (Step two). Meanwhile, the PCR-checked editor plasmid is transferred into the competent cells of *C. crescentus* through electroporation. The clones normally expressing *Sp*Cas9M are pre-screened by the indication of fluorescent sfGFP or antibiotic resistance (step three). Finally, the edited clones are confirmed by colony PCR and sequencing (step four). If necessary, the editor plasmid could be lost by cultivating the cells in antibiotics-free medium (step five). A complete genome editing workflow of CRISPR/*Sp*Cas9M-reporting system will be achieved in 5 days, and another 2-3 days may be needed when curing the plasmid.

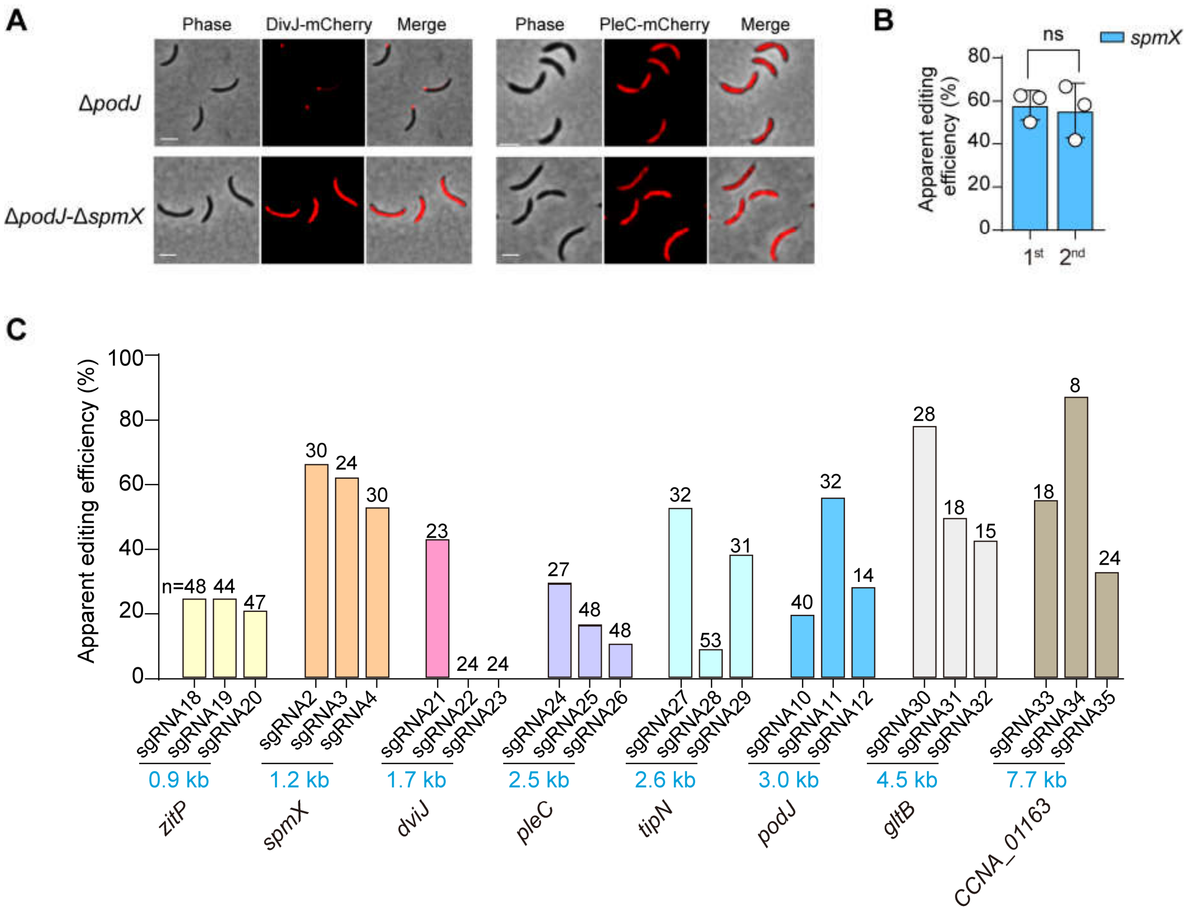


**Fig. S4. The CRISPR/*Sp*Cas9M-reporting system can be used for sequential and multiple-sites genome editing in *C. crescentus*.**

(*A*) The asymmetry of DivJ-mCherry and PleC-mCherry has completely lost in the double mutant Δ*podJ*-Δ*spmX*. The images of sequential knocking out of *podJ* and *spmX* are shown. All scale bars, 2 μm. (*B*) The apparent editing efficiency of the second-round editing in double mutation is comparable to that of the first-round editing in single mutation. The *spmX* gene was used as the editing target. All the data are presented as means ± SEM from 3 independent biological replicates. A total of 36 clones (*n*) were tested for each sample. ns, $p > 0.05$. (*C*) The CRISPR/*Sp*Cas9M-reporting system shows a well consistency in editing different gene targets. A series of genes with varying sizes (0.9-7.7 Kb) could be readily knocked out by the CRISPR/*Sp*Cas9M-reporting system, though distinct AAEs are shown. The numbers on the bars represent the tested clones in each sample.

### Table S1: The DNA sequence of *SpCas9*M

| ***Cas*** **gene** | **Sequence** |
|---|---|
| *SpCas9*M | atggacaagaagtactcgatcggcctggatatcgggacgaacagcgtcgggtgggccgtgatcacggacgaatacaaggtgcc<br>ctccaagaagttcaaggtcctggggaataccgaccggcactccatcaagaagaacctcatcggggccctgctctttgactccggg<br>gaaacggccgaagcgacgcggctgaagcggaccgcgcgtcgtcgctacacgcgccggaagaatcgcatctgttatctccagg<br>aaatcttcagcaacgagatggccaaggtggatgattccttcttccaccgcctggaggagtcctttctggtcgaagaggataagaag<br>catgagcgccacccgatctttgggaacatcgtcgacgaagtggcctaccacgaaaagtatcccacgatctaccacctgcgtaaga<br>agctcgtggactccaccgataaggcggatctgcggctgatctacctcgccctggcccacatgatcaagtttcggggccattttctga<br>tcgaaggcgatctgaatccggacaatagcgacgtcgataagctctttatccagctggtgcagacctacaatcaactcttcgaggag<br>aatccgatcaatgcctccggggtggatgccaaggcgatcctgagcgcgcgtctgtcgaagtcccgccgcctggagaacctgatc<br>gcccagctcccgggcgaaaagaagaacggctctttgggaatctgatcgccctcagcctggggctgaccccgaactttaagtcc<br>aatttcgacctcgcggaagatgccaagctccaactctcgaaggatacgtacgacgatgacctcgacaacctgctggcccaaatcg<br>gcgaccagtacgcggacctgttcctggcggccaagaacctgtcggacgccatcctcctgagcgacatcctccgtgtcaatacgg<br>aaatcacgaaggcgccgctgtccgccagcatgatcaagcggtatgatgagcatcatcaagacctcaccctgctgaaggcgctgg<br>tccggcaacagctgcccgagaagtacaaggagatctttttcgatcaaagcaagaacggctacgccgggtatatcgacgggggg<br>gcctcgcaagaagagttctacaagtttatcaagcccatcctggagaagatggatgggacggaggagctcctcgtcaagctcaacc<br>gggaagatctcctccgcaagcaacgcacctttgataatgggtccatcccgcatcaaatccacctcggcgagctccatgcgatcctg<br>cggcggcaagaggacttctacccgtttctcaaggacaaccgcgagaagatcgagaagatcctgacgttccgcatcccctattatgt<br>cggcccccctggcccgggggaattcgcggtttgcctggatgacccgcaagtccgaagaaacgatcacccccctggaattttgaaga<br>ggtggtggacaagggggcgagcgcgcagtcctttatcgaacggatgaccaatttcgacaagaacctccccaacgagaaggtgc<br>tcccgaagcattccctcctgtatgagtactttaccgtctacaacgagctcacgaaggtcaagtatgtcaccgagggcatgcgcaag<br>cccgccttcctctcgggggaacaaaagaaggccatcgtggacctgctcttcaagaccaaccgcaaggtcaccgtgaagcaactc<br>aaggaggactatttcaagaagatcgaatgcttcgatagcgtggagatctccggggtcgaagaccggtttaacgcctcgctcggca<br>cgtaccatgatctcctcaagatcatcaaggacaaggacttcctcgacaacgaggagaacgaagacatcctggaagacatcgtgct<br>gaccctgaccctcttcgaggatcgcgagatgatcgaggagcgtctcaagacgtacgcgcatctcttcgatgataaggtgatgaag<br>caactcaagcggcggcgttacacgggctgggggcgcctctcccgtaagctcatcaacggcatccgggacaagcaatcgggga<br>agaccatcctggacttcctgaagtcggacggctttgccaatcgcaattttatgcaactcatccacgacgattccctcaccttcaagga<br>agatatccagaaggcccaggtgtccgggcaaggggactccctgcatgaacatatcgccaacctggccgggagccccgcgatca<br>agaaggggatcctccagaccgtgaaggtggtcgatgagctcgtgaaggtcatgggccggcataagcccgaaaatatcgtcatcg<br>agatggcgcgcgagaaccagaccacccaaaagggccagaagaattcgcgggagcgcatgaagcggatcgaagaaggcatc<br>aaggaactcggcagccagatcctgaaggaacatccggtcgaaaacacgcagctccagaacgagaagctctacctctactatctc<br>caaaatggccgggatatgtacgtggaccaggagctcgacatcaatcggctcagcgactatgatgtggatcatatcgtcccccaatc<br>ctttctcaaggacgacagcatcgataacaaggtcctgacccggtcggacaagaatcgtggcaagagcgataacgtcccctccga<br>ggaggtcgtgaagaagatgaagaactattggcgccaactgctcaacgccaagctgatcacgcaacggaagttcgacaacctcac<br>caaggccgagcgcgggggcctctcggaactggacaaggcggggttcatcaagcgccagctcgtcgaaacccgtcaaatcacc<br>aagcacgtggcgcaaatcctggatagccgcatgaacaccaagtacgatgagaacgacaagctgatccgtgaagtgaaggtgat<br>caccctgaagtccaagctcgtgagcgatttccgcaaggatttccaattttataaggtgcgggaaatcaataattaccatcacgcgcat<br>gatgcgtatctcaacgccgtggtcggcacggccctgatcaagaagtatcccaagctggagtcggaattcgtctacggcgattaca<br>aggtgtatgacgtccgtaagatgatcgcgaagagcgagcaagaaatcgggaaggccacggcgaagtacttcttctacagcaata<br>tcatgaatttcttcaagacggagatcacgctggcgaatggcgaaatccgtaagcgcccgctgatcgaaacgaatggcgagaccg<br>gcgaaatcgtgtgggataagggggcgcgatttcgcgacggtccgcaaggtcctctcgatgccgcaggtcaacatcgtcaagaaga<br>ccgaagtccagaccgggggggttttccaaggagagcatcctgccgaagcgcaacagcgacaagctcatcgcccgcaagaagga<br>ctgggatcccaagaagtatggcgggttcgatagcccgaccgtggcgtactcggtgctggtggtcgcgaaggtcgaaaaggga |

| | |
|---|---|
| | agtccaagaagctcaagagcgtcaaggaactgctggggatcacgatcatggaacgctcctcctttgagaagaatcccatcgacttcctggaggcgaagggctataaggaggtgaagaaggatctcatcatcaagctgccgaagtattcgctcttcgaactcgaaaatgggcgcaagcgcatgctggcgtccgccggcgagctccaaaagggcaatgagctggccctgccctccaagtacgtcaattttctgtacctggcctcccactatgagaagctgaagggcagccccgaggataatgaacagaagcaactctttgtcgagcaacataagcattacctggatgaaatcatcgagcaaatcagcgaattttccaagcgggtgatcctcgccgatgccaacctggacaaggtcctcagcgcgtataacaagcaccgggacaagcccatccgtgaacaagcggagaacatcatccatctgttcacgctcaccaacctgggcgcccccgccgcgtttaagtattttgataccaccatcgatcgcaagcgttatacgtccaccaaggaagtcctcgatgccacgctgatccatcaaagcatcaccgggctctacgagacccggatcgatctctcccagctgggcggcgattga |

**Table S2: Putative neutral insertion sites in *C. crescentus* genome**

| No. | Start | Stop | Gap size | Convergent gap |
|---|---|---|---|---|
| 1 | 1996485 | 1996890 | 407 | Y |
| 2 | 2643951 | 2644381 | 432 | Y |
| 3 | 2655714 | 2656073 | 361 | Y |
| 4 | 2957896 | 2958322 | 428 | Y |
| 5 | 854686 | 855025 | 341 | Y |

Y, the corresponding annotation was passed.

### Table S3: Bacterial strains used in this study

| Strains | Description | Construction source or reference |
|---|---|---|
| ***C. crescentus*** | | |
| NA1000 | *C. crescentus* wild-type strain | Lucy Shapiro lab |
| JXS001 | *C. crescentus* NA1000 Δ*podJ* | One step knockout of *podJ* by pSJX019 in NA1000. This study. |
| JXS002 | *C. crescentus* NA1000 Δ*spmX* | One step knockout of *podJ* by pSJX016 in NA1000. This study. |
| JXS003 | *C. crescentus* NA1000 Δ*podJ*-Δ*spmX* | Second round knockout of *podJ* and *spmX* by pSJX019 and pSJX016 in NA1000. This study. |
| JXS004 | *C. crescentus* NA1000 Δ*zitP* | One step knockout of *zitP* by pSJX045 in NA1000. This study. |
| JXS005 | *C. crescentus* NA1000 Δ*divJ* | One step knockout of *divJ* by pSJX048 in NA1000. This study. |
| JXS006 | *C. crescentus* NA1000 Δ*pleC* | One step knockout of *pleC* by pSJX051 in NA1000. This study. |
| JXS007 | *C. crescentus* NA1000 Δ*tipN* | One step knockout of *tipN* by pSJX054 in NA1000. This study. |
| JXS008 | *C. crescentus* NA1000 Δ*gltB* | One step knockout of *gltB* by pSJX057 in NA1000. This study. |
| JXS009 | *C. crescentus* NA1000 Δ*CCNA_01163* | One step knockout of *CCNA_01163* by pSJX060 in NA1000. This study. |
| JXS010 | $Chl^R$; *C. crescentus* NA1000, *pBVMCS6-*$P_{van}$*-pleC-mCherry* | This study. |
| JXS011 | $Chl^R$; *C. crescentus* NA1000 Δ*podJ*, *pBVMCS6-*$P_{van}$*-pleC-mCherry* | This study. |
| JXS012 | $Chl^R$; *C. crescentus* NA1000, *pBVMCS6-*$P_{van}$*-divJ-mCherry* | This study. |
| JXS013 | $Chl^R$; *C. crescentus* NA1000 Δ*podJ*, *pBVMCS6-*$P_{van}$*-divJ-mCherry* | This study. |
| JXS014 | $Chl^R$; *C. crescentus* NA1000 Δ*podJ*-Δ*spmX*, *pBVMCS6-*$P_{van}$*-pleC-mCherry* | This study. |
| JXS015 | $Chl^R$; *C. crescentus* NA1000 Δ*podJ*-Δ*spmX*, *pBVMCS6-*$P_{van}$*-divJ-mCherry* | This study. |
| JXS016 | *C. crescentus* NA1000, *NIS1::*$P_{cat}$*-mCherry* | One step knock-in of $P_{cat}$*-mCherry* at NIS1 site by pSJX071 in NA1000. This study. |
| JXS017 | *C. crescentus* NA1000, *NIS2::*$P_{cat}$*-mCherry* | One step knock-in of $P_{cat}$*-mCherry* at NIS2 site by pSJX072 in NA1000. This study. |
| JXS018 | *C. crescentus* NA1000 Δ*spmX::mCherry* | One step substitution of *spmX* with *mCherry* by pSJX072 in NA1000. This study. |
| ***A. fabrum*** | | |
| C58 | *A. fabrum* wild-type strain | Addgene |
| JXS019 | *A. fabrum* C58 Δ*tdk* | One step knockout of *tdk* by pSJX073 in *A. fabrum* C58. This study. |
| ***S. meliloti*** | | |
| 1021 | *S. meliloti* wild-type strain | Zhongjun qin lab |
| JXS020 | *S. meliloti* 1021 Δ*tdk* | One step knockout of *tdk* by pSJX076 in *S. meliloti* 1021. This study. |
| ***E. coli*** | | |

| | | |
|---|---|---|
| DH5α | Bacterial cloning strain | Novagen |
| WM6026 | Bacterial conjugation | Yi Song lab |

[a]Abbreviations: Chl, Chloramphenicol; R, resistance.

### Table S4: Plasmids used in this study

| Plasmid | Plasmid information[a] | Source or reference |
|---|---|---|
| pSJX001 | $Kan^R$; pBVMCS2-sgRNA2-Δ*spmX* (H-arms)-$P_{van}$-*SpCas9* | This study |
| pSJX002 | $Kan^R$; pBVMCS2-Δ*spmX* (H-arms)-$P_{van}$-*SpCas9* | This study |
| pSJX003 | $Kan^R$; pBVMCS2-sgRNA1-Δ*spmX* (H-arms)-$P_{van}$-*SpCas9* | This study |
| pSJX004 | $Kan^R$; pBVMCS2-sgRNA2-Δ*spmX* (H-arms)-$P_{van}$ | This study |
| pSJX005 | $Kan^R$; pBVMCS2-sgRNA2-Δ*spmX* (H-arms)-$P_{van}$-*SpCas9*M | This study |
| pSJX006 | $Kan^R$; pBVMCS2-sgRNA7-Δ*spmX*(H-arms)-$P_{van}$-*FnCas12a*M | This study |
| pSJX007 | $Kan^R$; pBVMCS2-sgRNA8-Δ*spmX* (H-arms)-$P_{van}$-*Sth1Cas9*M | This study |
| pSJX008 | $Kan^R$; pBVMCS2-sgRNA9-Δ*spmX* (H-arms)-$P_{van}$-*Sth3Cas9*M | This study |
| pSJX009 | $Kan^R$; pBVMCS2-sgRNA10-Δ*podJ* (H-arms)-$P_{van}$ | This study |
| pSJX010 | $Kan^R$; pBVMCS2-sgRNA10-Δ*podJ* (H-arms)-$P_{van}$-*SpCas9* | This study |
| pSJX011 | $Kan^R$; pBVMCS2-sgRNA10-Δ*podJ* (H-arms)-$P_{van}$-*SpCas9*M | This study |
| pSJX012 | $Kan^R$; pBVMCS2-sgRNA13-Δ*podJ*(H-arms)-$P_{van}$-*FnCas12a*M | This study |
| pSJX013 | $Kan^R$; pBVMCS2-sgRNA14-Δ*podJ* (H-arms)-$P_{van}$-*Sth1Cas9*M | This study |
| pSJX014 | $Kan^R$; pBVMCS2-sgRNA15-Δ*podJ* (H-arms)-$P_{van}$-*Sth3Cas9*M | This study |
| pSJX015 | $Kan^R$; pBXMCS2-sgRNA2-Δ*spmX* (H-arms)-$P_{xyl}$-*SpCas9*M | This study |
| pSJX016 | $Kan^R$; pBVMCS2-sgRNA2-Δ*spmX* (H-arms)-$P_{van}$-*SpCas9*M-*sfgfp* | This study |
| pSJX017 | $Kan^R$; pBVMCS2-sgRNA3-Δ*spmX* (H-arms)-$P_{van}$-*SpCas9*M-*sfgfp* | This study |
| pSJX018 | $Kan^R$; pBVMCS2-sgRNA4-Δ*spmX* (H-arms)-$P_{van}$-*SpCas9*M-*sfgfp* | This study |
| pSJX019 | $Kan^R$; pBVMCS2-sgRNA10-Δ*podJ* (H-arms)-$P_{van}$-*SpCas9*M-*sfgfp* | This study |
| pSJX020 | $Kan^R$; pBVMCS2-sgRNA11-Δ*podJ* (H-arms)-$P_{van}$-*SpCas9*M-*sfgfp* | This study |
| pSJX021 | $Kan^R$; pBVMCS2-sgRNA12-Δ*podJ* (H-arms)-$P_{van}$-*SpCas9*M-*sfgfp* | This study |
| pSJX022 | $Kan^R$; pBVMCS2-sgRNA2-Δ*spmX* (H-arms)-$P_{van}$-*SpCas9*M-*cat* | This study |
| pSJX023 | $Kan^R$; pBXMCS2-sgRNA2-Δ*spmX* (H-arms)-$P_{xyl}$-*SpCas9*M-*sfgfp* | This study |
| pSJX024 | $Kan^R$; pBXMCS2-sgRNA2-Δ*spmX* (H-arms)-$P_{xyl}$-*SpCas9*M-*cat* | This study |
| pSJX025 | $Spec^R$; pBVMCS1-sgRNA2-Δ*spmX* (H-arms)-$P_{van}$-*SpCas9*M-*sfgfp* | This study |
| pSJX026 | $Spec^R$; pBVMCS1-sgRNA2-Δ*spmX* (H-arms)-$P_{van}$-*SpCas9*M-*cat* | This study |
| pSJX027 | $Spec^R$; pBXMCS1-sgRNA2-Δ*spmX* (H-arms)-$P_{xyl}$-*SpCas9*M-*sfgfp* | This study |
| pSJX028 | $Spec^R$; pBXMCS1-sgRNA2-Δ*spmX* (H-arms)-$P_{xyl}$-*SpCas9*M-*cat* | This study |
| pSJX029 | $Kan^R$; pBVMCS2-sgRNA3-Δ*spmX* (H-arms)-$P_{van}$-*SpCas9*M | This study |
| pSJX030 | $Kan^R$; pBVMCS2-sgRNA4-Δ*spmX* (H-arms)-$P_{van}$-*SpCas9*M | This study |
| pSJX031 | $Kan^R$; pBVMCS2-sgRNA3-Δ*spmX* (H-arms)-$P_{van}$-*SpCas9*M-*cat* | This study |
| pSJX032 | $Kan^R$; pBVMCS2-sgRNA4-Δ*spmX* (H-arms)-$P_{van}$-*SpCas9*M-*cat* | This study |
| pSJX033 | $Kan^R$; pBVMCS2-sgRNA2-Δ*spmX*(H-arms-0bp)-$P_{van}$-*SpCas9*M | This study |
| pSJX034 | $Kan^R$; pBVMCS2-sgRNA2-Δ*spmX* (H-arms-50bp)-$P_{van}$-*SpCas9*M | This study |
| pSJX035 | $Kan^R$; pBVMCS2-sgRNA2-Δ*spmX* (H-arms-200bp)-$P_{van}$-*SpCas9*M | This study |
| pSJX036 | $Kan^R$; pBVMCS2-sgRNA2-Δ*spmX* (H-arms-500bp)-$P_{van}$-*SpCas9*M | This study |
| pSJX037 | $Kan^R$; pBVMCS2-sgRNA2-Δ*spmX* (H-arms-1000bp)-$P_{van}$-*SpCas9*M | This study |
| pSJX038 | $Kan^R$; pBMCS2-sgRNA2-Δ*spmX* (H-arms)-*SpCas9*M (no promoter) | This study |
| pSJX039 | $Kan^R$; pBMCS2-sgRNA10-Δ*podJ* (H-arms)-*SpCas9*M (no promoter) | This study |
| pSJX040 | $Kan^R$; pBXMCS2-sgRNA10-Δ*podJ* (H-arms)-$P_{xyl}$-*SpCas9*M | This study |

| | | |
|---|---|---|
| pSJX041 | Kan$^{R}$; pBVMCS2-sgRNA5-Δ*spmX* (H-arms)-P$_{van}$-*SpCas9*M | This study |
| pSJX042 | Kan$^{R}$; pBVMCS2-sgRNA6-Δ*spmX* (H-arms)-P$_{van}$-*SpCas9*M | This study |
| pSJX043 | Chl$^{R}$; pBVMCS-6-P$_{van}$-*divJ-mCherry* | This study |
| pWT194 | Chl$^{R}$; pBVMCS-6-P$_{van}$-*pleC-mCherry* | (12) |
| pSJX045 | Kan$^{R}$; pBVMCS2-sgRNA18-*zitP* (H-arms)-P$_{van}$-*SpCas9*M-*sfgfp* | This study |
| pSJX046 | Kan$^{R}$; pBVMCS2-sgRNA19-*zitP* (H-arms)-P$_{van}$-*SpCas9*M-*sfgfp* | This study |
| pSJX047 | Kan$^{R}$; pBVMCS2-sgRNA20-*zitP* (H-arms)-P$_{van}$-*SpCas9*M-*sfgfp* | This study |
| pSJX048 | Kan$^{R}$; pBVMCS2-sgRNA21-*divJ* (H-arms)-P$_{van}$-*SpCas9*M-*sfgfp* | This study |
| pSJX049 | Kan$^{R}$; pBVMCS2-sgRNA22-*divJ* (H-arms)-P$_{van}$-*SpCas9*M-*sfgfp* | This study |
| pSJX050 | Kan$^{R}$; pBVMCS2-sgRNA23-*divJ* (H-arms)-P$_{van}$-*SpCas9*M-*sfgfp* | This study |
| pSJX051 | Kan$^{R}$; pBVMCS2-sgRNA24-*pleC* (H-arms)-P$_{van}$-*SpCas9*M-*sfgfp* | This study |
| pSJX052 | Kan$^{R}$; pBVMCS2-sgRNA25-*pleC* (H-arms)-P$_{van}$-*SpCas9*M-*sfgfp* | This study |
| pSJX053 | Kan$^{R}$; pBVMCS2-sgRNA26-*pleC* (H-arms)-P$_{van}$-*SpCas9*M-*sfgfp* | This study |
| pSJX054 | Kan$^{R}$; pBVMCS2-sgRNA27-*tipN* (H-arms)-P$_{van}$-*SpCas9*M-*sfgfp* | This study |
| pSJX055 | Kan$^{R}$; pBVMCS2-sgRNA28-*tipN* (H-arms)-P$_{van}$-*SpCas9*M-*sfgfp* | This study |
| pSJX056 | Kan$^{R}$; pBVMCS2-sgRNA29-*tipN* (H-arms)-P$_{van}$-*SpCas9*M-*sfgfp* | This study |
| pSJX057 | Kan$^{R}$; pBVMCS2-sgRNA30-*gltB* (H-arms)-P$_{van}$-*SpCas9*M-*sfgfp* | This study |
| pSJX058 | Kan$^{R}$; pBVMCS2-sgRNA31-*gltB* (H-arms)-P$_{van}$-*SpCas9*M-*sfgfp* | This study |
| pSJX059 | Kan$^{R}$; pBVMCS2-sgRNA32-*gltB* (H-arms)-P$_{van}$-*SpCas9*M-*sfgfp* | This study |
| pSJX060 | Kan$^{R}$; pBVMCS2-sgRNA33-*CCNA_01163* (H-arms)-P$_{van}$-*SpCas9*M-*sfgfp* | This study |
| pSJX061 | Kan$^{R}$; pBVMCS2-sgRNA34-*CCNA_01163* (H-arms)-P$_{van}$-*SpCas9*M-*sfgfp* | This study |
| pSJX062 | Kan$^{R}$; pBVMCS2-sgRNA35-*CCNA_01163* (H-arms)-P$_{van}$-*SpCas9*M-*sfgfp* | This study |
| pSJX063 | Kan$^{R}$; pBVMCS2-sgRNA2-Δ*spmX* (Larm-*mCherry*-Rarm)-P$_{van}$-*SpCas9*M-*cat* | This study |
| pSJX064 | Kan$^{R}$; pBVMCS2-sgRNA3-Δ*spmX* (Larm-*mCherry*-Rarm)-P$_{van}$-*SpCas9*M-*cat* | This study |
| pSJX065 | Kan$^{R}$; pBVMCS2-sgRNA16-NIS1 (Larm-P$_{cat}$-*mCherry*-Rarm)-P$_{van}$-*SpCas9*M-*cat* | This study |
| pSJX066 | Kan$^{R}$; pBVMCS2-sgRNA17-NIS2 (Larm-P$_{cat}$-*mCherry*-Rarm)-P$_{van}$-*SpCas9*M-*cat* | This study |
| pSJX067 | Kan$^{R}$; pBVMCS2-sgRNA36-*tdk* (H-arms)-P$_{van}$-*SpCas9*M-*sfgfp* | This study |
| pSJX068 | Kan$^{R}$; pBVMCS2-sgRNA37-*tdk* (H-arms)-P$_{van}$-*SpCas9*M-*sfgfp* | This study |
| pSJX069 | Kan$^{R}$; pBVMCS2-sgRNA38-*tdk* (H-arms)-P$_{van}$-*SpCas9*M-*sfgfp* | This study |
| pSJX070 | Gent$^{R}$; pK18mob-P$_{lac}$-*SpCas9*M-*sfgfp*-sgRNA39-*tdk* (H-arms) | This study |
| pSJX071 | Gent$^{R}$; pK18mob-P$_{lac}$-*SpCas9*M-*sfgfp*-sgRNA40-*tdk* (H-arms) | This study |
| pSJX072 | Gent$^{R}$; pK18mob-P$_{lac}$-*SpCas9*M-*sfgfp*-sgRNA41-*tdk* (H-arms) | This study |

[a]Abbreviations: Kan, kanamycin; Chl, chloramphenicol; Spec, Spectinomycin; Gent, Gentamicin; R, resistance.

### Table S5: sgRNAs used in this study

| sgRNA | Sequences | Description |
|---|---|---|
| sgRNA1 | GAAGAGCGAGATCTCGAGCT | NT-sgRNA |
| sgRNA2 | TCTCTTGATGAGATCGACGG | Target *spmX* for *Sp*Cas9 and *Sp*Cas9M |
| sgRNA3 | GAAGACTCTGTTCCTCACGC | Target *spmX* for *Sp*Cas9M |
| sgRNA4 | ATCTCATCAAGAGATTCGAA | Target *spmX* for *Sp*Cas9M |
| sgRNA5 | GAGGAACAGAGTCTTCTCGG | Target *spmX* for *Sp*Cas9M |
| sgRNA6 | CGTGAGGAACAGAGTCTTCT | Target *spmX* for *Sp*Cas9M |
| sgRNA7 | GGAGGAAGTTGTCGAGGCCGATATT | Target *spmX* for *Fn*Cas12aM |
| sgRNA8 | ACGGTTGCGTCCTCGCTGGT | Target *spmX* for *Sth*1Cas9M |
| sgRNA9 | CTACTCTTCGTCGCTCACAT | Target *spmX* for *Sth*3Cas9M |
| sgRNA10 | CGCCGATAGCGTTCAAGCCC | Target *podJ* for *Sp*Cas9 and *Sp*Cas9M |
| sgRNA11 | CTTCGAGAGGTAGAACTGCG | Target *podJ* for *Sp*Cas9M |
| sgRNA12 | CATGCGATCGAAACGTCCGT | Target *podJ* for *Sp*Cas9M |
| sgRNA13 | GCCGAGCTTGCGCTCCATGCGATC | Target *podJ* for *Fn*Cas12aM |
| sgRNA14 | AGCGCACGCTCCGCATGCTG | Target *podJ* for *Sth*1Cas9M |
| sgRNA15 | GATCGACAGCCGTCTGTCCG | Target *podJ* for *Sth*3Cas9M |
| sgRNA16 | CATCGAGGCAGAAGATGCAC | Target NIS1 for *Sp*Cas9M |
| sgRNA17 | GGTCAACGCAGCGCCGACCA | Target NIS2 for *Sp*Cas9M |
| sgRNA18 | CGGCGAGATGACCTCATGCT | Target *zitP* for *Sp*Cas9M |
| sgRNA19 | GGCTTCAGGCTTCTTCAGCG | Target *zitP* for *Sp*Cas9M |
| sgRNA20 | TTGGGCCTGGTGATCGACGC | Target *zitP* for *Sp*Cas9M |
| sgRNA21 | GCTGAGCTGATCCACGAGAG | Target *divJ* for *Sp*Cas9M |
| sgRNA22 | GGTCTGGAGACCTTGCTCGA | Target *divJ* for *Sp*Cas9M |
| sgRNA23 | ACCGCCGGAGTTCAGCTGCG | Target *divJ* for *Sp*Cas9M |
| sgRNA24 | GCGCAAGTACGAGACCGAGA | Target *pleC* for *Sp*Cas9M |
| sgRNA25 | GCATCTCGAAGACGTCGCCG | Target *pleC* for *Sp*Cas9M |
| sgRNA26 | ATCCTCGATTGAGAGCGCAA | Target *pleC* for *Sp*Cas9M |
| sgRNA27 | CGAACCCCGAAGATCCAGGG | Target *tipN* for *Sp*Cas9M |
| sgRNA28 | TCTCGTTCAGCTGATCGACA | Target *tipN* for *Sp*Cas9M |
| sgRNA29 | GCTCGAAGAAGAGCTGAAGG | Target *tipN* for *Sp*Cas9M |
| sgRNA30 | AAACCGGTGATCCAGCCGGG | Target *gltB* for *Sp*Cas9M |
| sgRNA31 | GCTCTATTCCTACTGCAACG | Target *gltB* for *Sp*Cas9M |
| sgRNA32 | GTTGGGCGAGAAGAACGTCG | Target *gltB* for *Sp*Cas9M |
| sgRNA33 | GGGTGGCATTAAGAGCCGAG | Target *CCNA_01163* for *Sp*Cas9M |
| sgRNA34 | GATCTGGTCAGCGGTCAGGG | Target *CCNA_01163* for *Sp*Cas9M |
| sgRNA35 | CAGGGCCCCGATCTGCACTG | Target *CCNA_01163* for *Sp*Cas9M |
| sgRNA36 | GAAGGCGCCCAGATCGAAGT | Target *atu-tdk* for *Sp*Cas9M |
| sgRNA37 | CAGATCGAAGTTGGCGGCAA | Target *atu-tdk* for *Sp*Cas9M |
| sgRNA38 | AGATCGAAGTTGGCGGCAAC | Target *atu-tdk* for *Sp*Cas9M |
| sgRNA39 | AGCTATGCGACGATGAATGC | Target *sme-tdk* for *Sp*Cas9M |
| sgRNA40 | GGTCGCATCGCGTCTCGCAT | Target *sme-tdk* for *Sp*Cas9M |

| | | |
|---|---|---|
| sgRNA41 | TCTCCTATTGCCGTCGCCAT | Target *sme-tdk* for *Sp*Cas9M |

**Table S6: Oligonucleotides used in this study**

| Primer | Sequence | Name of plasmids |
|---|---|---|
| SJX0299 | tagctgtcaagcgtggagcattcgcgcc | pSJX001 |
| SJX0300 | tgctccacgcttgacagctagctcagtcctaggtataatact | |
| SJX0326 | ttgtttgtcgggctggactctagccgac | |
| SJX0325 | gagtccagcccgacaaacaacagataaaacgaaaggccc | |
| SJX1219 | ggaaacgcatatggataagaaatactcaataggcttagatatcgg | |
| SJX1222 | ccgttttcattcagtcacctcctagctgactcaaa | |
| SJX1221 | aggtgactgaatgaaaacgggccccccc | |
| SJX1220 | tcttatccatatgcgtttcctcgcatcg | |
| SJX0514 | tgcttttttacaatcacccgatcgccc | |
| SJX0517 | ggcaagaaaccgcctagattgacccagtttgag | |
| SJX0516 | aatctaggcggtttcttgcctcgccacc | |
| SJX0519 | gctgggcgcgccagcttggaccgatcgt | |
| SJX0518 | tccaagctggcgcgcccagcggcgcgga | |
| SJX0515 | gggtgattgtaaaaaaagcaccgactcggtg | |
| SJX0652 | tctcttgatgagatcgacgggttttagagctagaaatagcaagttaaaataaggctagtc | |
| SJX0653 | ccgtcgatctcatcaagagaactagtattatacctaggactgagctagctg | |
| SJX0301 | gtcctaggtataatactagtgttttagagctagaaatagcaagttaaaataaggctagt | pSJX002 |
| SJX0302 | gctatttctagctctaaaacactagtattatacctaggactgagctagctg | |
| SJX1525 | gaagagcgagatctcgagctgttttagagctagaaatagcaagttaaaataaggctagt | pSJX003 |
| SJX1523 | agctcgagatctcgctcttcactagtattatacctaggactgagctagctg | |
| SJX1520 | ggaaacgcatatgaaaacgggccccccc | pSJX004 |
| SJX1521 | ccgttttcatatgcgtttcctcgcatcgtg | |
| SJX0295 | ggaaacgcatatggacaagaagtactcgatcgg | pSJX005 |
| SJX0296 | ccgttttcattcaatcgccgcccagctg | |
| SJX0297 | cggcgattgaatgaaaacgggccccccc | |
| SJX0298 | tcttgtccatatgcgtttcctcgcatcg | |
| SJX1146 | ggaaacgcatatgagcatctaccaggagttcgt | pSJX006 |
| SJX1147 | agatgctcatatgcgtttcctcgcatcgt | |
| SJX1148 | ccgttttcattcaattgttccgattctgcacaaattc | |
| SJX1149 | gaacaattgaatgaaaacgggccccccc | |
| SJX1233 | ggaggaagttgtcgaggccgatattatctacaacagtagaaattccacaatcacccgatcg ccc | |
| SJX1234 | cggcctcgacaacttcctccactagtattatacctaggactgagctagct | |
| SJX1269 | acggttgcgtcctcgctggtgtttttgtactcgaaagaagctacaaagataagg | pSJX007 |
| SJX1270 | accagcgaggacgcaaccgtactagtattatacctaggactgagctagctg | |
| SJX1201 | gtgtttttttacaatcacccgatcgcccgc | |
| SJX1202 | gggtgattgtaaaaaaacaccctgccataaaatgacagggt | |
| SJX1203 | ggaaacgcatatgtcggacctggtcctgg | |
| SJX1204 | ggtccgacatatgcgtttcctcgcatcg | |
| SJX0065 | agccgaagctggacttctgaatgaaaacgggcccccc | |
| SJX0068 | gagggggggcccgttttcattcagaagtccagcttcggct | |
| SJX1248 | ctactcttcgtcgctcacatgttttagagctgtgaaaacagcgagt | pSJX008 |
| SJX1249 | atgtgagcgacgaagagtagactagtattatacctaggactgagctagct | |

| | | |
|---|---|---|
| SJX1268 | gtgtttttttacaatcacccgatcgcccgc | |
| SJX1269 | gggtgattgtaaaaaaacaccgaatcggtgccacc | |
| SJX1158 | ggaaacgcatatgaccaagccgtactcgatcg | |
| SJX1159 | gcttggtcatatgcgtttcctcgcatcg | |
| SJX0069 | ccaagctgggcgagggctgaatgaaaacgggcccccc | |
| SJX0072 | gagggggggcccgttttcattcagccctcgcccagc | |
| SJX0715 | cgccgatagcgttcaagcccgttttagagctagaaatagcaagttaaaataaggc | pSJX010 |
| SJX0716 | gggcttgaacgctatcggcgactagtattatacctaggactgagctagct | |
| SJX1217 | tgcttttttttgctgttggcctccgggg | |
| SJX1218 | gccaacagcaaaaaaaagcaccgactcggtg | |
| SJX0312 | atcgattcgcacgcctcgcgacctcgcg | |
| SJX0311 | cgcgaggcgtgcgaatcgatctccccgcac | |
| SJX1194 | cacccggcaacgcgcccagcggcgcgga | |
| SJX1193 | gctgggcgcgttgccgggtgacgtgatc | |
| SJX1520 | ggaaacgcatatgaaaacgggccccccc | pSJX009 |
| SJX1521 | ccgttttcatatgcgtttcctcgcatcgtg | |
| SJX0295 | ggaaacgcatatggacaagaagtactcgatcgg | pSJX011 |
| SJX0296 | ccgttttcattcaatcgccgcccagctg | |
| SJX0297 | cggcgattgaatgaaaacgggccccccc | |
| SJX0298 | tcttgtccatatgcgtttcctcgcatcg | |
| SJX1191 | agcttgcgctccatgcgatcatctacaacagtagaaattcctgctgttggcctccgggg | pSJX012 |
| SJX1192 | gatcgcatggagcgcaagctcggcactagtattatacctaggactgagctagct | |
| SJX1146 | ggaaacgcatatgagcatctaccaggagttcgt | |
| SJX1147 | agatgctcatatgcgtttcctcgcatcgt | |
| SJX1148 | ccgttttcattcaattgttccgattctgcacaaattc | |
| SJX1149 | gaacaattgaatgaaaacgggccccccc | |
| SJX1193 | agcgcacgctccgcatgctggtttttgtactcgaaagaagctacaaagataagg | pSJX013 |
| SJX1194 | cagcatgcggagcgtgcgctactagtattatacctaggactgagctagct | |
| SJX1215 | gtgtttttttgctgttggcctccgggg | |
| SJX1216 | gccaacagcaaaaaaaacaccctgccataaaatgacag | |
| SJX1203 | ggaaacgcatatgtcggacctggtcctgg | |
| SJX1204 | ggtccgacatatgcgtttcctcgcatcg | |
| SJX1195 | gatcgacagccgtctgtccggtttttgtactcgaaagaagctacaaagataagg | pSJX014 |
| SJX1196 | cggacagacggctgtcgatcactagtattatacctaggactgagctagct | |
| SJX1215 | gtgtttttttgctgttggcctccgggg | |
| SJX1216 | gccaacagcaaaaaaaacaccctgccataaaatgacag | |
| SJX1158 | ggaaacgcatatgaccaagccgtactcgatcg | |
| SJX1159 | gcttggtcatatgcgtttcctcgcatcg | |
| SJX1326 | tgttgtttgtcgctgcagccagccgtggtc | pSJX015 |
| SJX1327 | ggctggctgcagcgacaaacaacagataaaacgaaaggc | |
| SJX1328 | tggggagacgaccatatggacaagaagtactcgatcgg | |
| SJX1329 | gtacttcttgtccatatggtcgtctccccaaaa | |
| SJX1577 | ccgaccggtgatcgccgcccagctggga | pSJX016 |
| SJX1580 | gggcggcgatcaccggtcggccaccatgcgtaaaggcgaagagctg | |
| SJX1581 | gtacaaatgaatgaaaacgggcccccccctc | |
| SJX1582 | ccgttttcattcatttgtacagttcatccataccatgcgt | |
| SJX1737 | gaagactctgttcctcacgcgttttagagctagaaatagcaagttaaaataaggctag | pSJX017 |
| SJX1738 | gcgtgaggaacagagtcttcactagtattatacctaggactgagctagctg | |

| | | |
|---|---|---|
| SJX1739 | atctcatcaagagattcgaagttttagagctagaaatagcaagttaaaataaggctag | pSJX018 |
| SJX1740 | ttcgaatctcttgatgagatactagtattatacctaggactgagctagctg | |
| SJX1577 | ccgaccggtgatcgccgcccagctggga | pSJX019 |
| SJX1580 | gggcggcgatcaccggtcggccaccatgcgtaaaggcgaagagctg | |
| SJX1581 | gtacaaatgaatgaaaacgggcccccctc | |
| SJX1582 | ccgttttcattcatttgtacagttcatccataccatgcgt | |
| SJX1733 | cttcgagaggtagaactgcggttttagagctagaaatagcaagttaaaataaggctag | pSJX020 |
| SJX1734 | cgcagttctacctctcgaagactagtattatacctaggactgagctagct | |
| SJX1735 | catgcgatcgaaacgtccgtgttttagagctagaaatagcaagttaaaataaggctag | pSJX021 |
| SJX1736 | acggacgtttcgatcgcatgactagtattatacctaggactgagctagct | |
| SJX1874 | accggtcggccaccatggagaaaaaaatcactggatataccacc | pSJX022 |
| SJX1577 | ccgaccggtgatcgccgcccagctggga | |
| SJX1831 | gcccgttttcatttacgccccgccctgcca | |
| SJX1832 | gcggggcgtaaatgaaaacgggcccccccct | |
| SJX1577 | ccgaccggtgatcgccgcccagctggga | pSJX023 |
| SJX1580 | gggcggcgatcaccggtcggccaccatgcgtaaaggcgaagagctg | |
| SJX1581 | gtacaaatgaatgaaaacgggcccccctc | |
| SJX1582 | ccgttttcattcatttgtacagttcatccataccatgcgt | |
| SJX1874 | accggtcggccaccatggagaaaaaaatcactggatataccacc | pSJX024 |
| SJX1577 | ccgaccggtgatcgccgcccagctggga | |
| SJX1831 | gcccgttttcatttacgccccgccctgcca | |
| SJX1832 | gcggggcgtaaatgaaaacgggcccccccct | |
| SJX2393 | tgctccacgcttgacagctagctcagtcctaggtataa | pSJX025 |
| SJX2394 | tagctgtcaagcgtggagcattcgcgcc | |
| SJX1581 | ccgttttcattcatttgtacagttcatccataccatgcgt | |
| SJX1582 | ccgttttcattcatttgtacagttcatccataccatgcgt | |
| SJX2393 | tgctccacgcttgacagctagctcagtcctaggtataa | pSJX026 |
| SJX2394 | tagctgtcaagcgtggagcattcgcgcc | |
| SJX1831 | gcccgttttcatttacgccccgccctgcca | |
| SJX1832 | gcggggcgtaaatgaaaacgggcccccccct | |
| SJX2393 | tgctccacgcttgacagctagctcagtcctaggtataa | pSJX027 |
| SJX2394 | tagctgtcaagcgtggagcattcgcgcc | |
| SJX1581 | ccgttttcattcatttgtacagttcatccataccatgcgt | |
| SJX1582 | ccgttttcattcatttgtacagttcatccataccatgcgt | |
| SJX2393 | tgctccacgcttgacagctagctcagtcctaggtataa | pSJX028 |
| SJX2394 | tagctgtcaagcgtggagcattcgcgcc | |
| SJX1831 | gcccgttttcatttacgccccgccctgcca | |
| SJX1832 | gcggggcgtaaatgaaaacgggcccccccct | |
| SJX1737 | gaagactctgttcctcacgcgttttagagctagaaatagcaagttaaaataaggctag | pSJX029 |
| SJX1738 | gcgtgaggaacagagtcttcactagtattatacctaggactgagctagctg | |
| SJX1739 | atctcatcaagagattcgaagttttagagctagaaatagcaagttaaaataaggctag | pSJX030 |
| SJX1740 | ttcgaatctcttgatgagatactagtattatacctaggactgagctagctg | |
| SJX1737 | gaagactctgttcctcacgcgttttagagctagaaatagcaagttaaaataaggctag | pSJX031 |
| SJX1738 | gcgtgaggaacagagtcttcactagtattatacctaggactgagctagctg | |
| SJX1739 | atctcatcaagagattcgaagttttagagctagaaatagcaagttaaaataaggctag | pSJX032 |
| SJX1740 | ttcgaatctcttgatgagatactagtattatacctaggactgagctagctg | |
| SJX1336 | tgcttttttcgcgcccagcggcgcggatc | pSJX033 |
| SJX1337 | gctgggcgcgaaaaaaagcaccgactcggtgc | |

| | | |
|---|---|---|
| SJX1350 | cggtgcttttttgggacaggcgaacgcatc | pSJX034 |
| SJX1351 | cgttcgcctgtcccaaaaaaagcaccgactcggt | |
| SJX1352 | cgctggtccccgcgcccagcggcgcgga | |
| SJX1353 | gctgggcgcggggaccagcgggcgttct | |
| SJX1346 | cggtgcttttttacccggcggtgtcgtcct | pSJX035 |
| SJX1347 | gacaccgccgggtaaaaaaagcaccgactcggtgcc | |
| SJX1348 | ggaaggaaaccgcgcccagcggcgcgga | |
| SJX1349 | gctgggcgcggtttccttcccgaacgcc | |
| SJX1342 | cggtgcttttttttctcgtccctgggcggca | pSJX036 |
| SJX1343 | gcccagggacgagaaaaaaaagcaccgactcggtg | |
| SJX1344 | cccactccaacgcgcccagcggcgcgga | |
| SJX1345 | gctgggcgcgttggagtgggcgccgtag | |
| SJX1338 | gtcggtgcttttttgacgatcgaccatgtccgc | pSJX037 |
| SJX1339 | acatggtcgatcgtcaaaaaaagcaccgactcggt | |
| SJX1340 | ctcgatgtcgcgcgcccagcggcgcgga | |
| SJX1341 | gctgggcgcgcgacatcgaggcaagcct | |
| SJX1330 | ttgtttgtcgatggacaagaagtactcgatcggc | pSJX038 |
| SJX1331 | tcttgtccatcgacaaacaacagataaaacgaaaggcc | |
| SJX1330 | ttgtttgtcgatggacaagaagtactcgatcggc | pSJX039 |
| SJX1331 | tcttgtccatcgacaaacaacagataaaacgaaaggcc | |
| SJX1326 | tgttgtttgtcgctgcagccagccgtggtc | pSJX040 |
| SJX1327 | ggctggctgcagcgacaaacaacagataaaacgaaaggc | |
| SJX1328 | tggggagacgaccatatggacaagaagtactcgatcgg | |
| SJX1329 | gtacttcttgtccatatggtcgtctccccaaaa | |
| SJX0650 | gaggaacagagtcttctcgggttttagagctagaaatagcaagttaaaataaggctagtc | pSJX041 |
| SJX0651 | ccgagaagactctgttcctcactagtattatacctaggactgagctagctg | |
| SJX1332 | cgtgaggaacagagtcttctgttttagagctagaaatagcaagttaaaataaggctagtc | pSJX042 |
| SJX1333 | gaagactctgttcctcacgactagtattatacctaggactgagctagctg | |
| SJX1257 | cgaggaaacgttggaattcgaaacgcttccagaccc | pSJX043 |
| SJX1258 | cgaattccaacgtttcctcgcatcgtgg | |
| SJX1259 | tgcgccgcgccaccggtcggccaccatgg | |
| SJX1260 | ccgaccggtggcgcggcgcaaaggcgat | |
| SJX1597 | ggtgcttttttgttggcggcggcgaagcc | pSJX045 |
| SJX1598 | gccgccgccaacaaaaaaagcaccgactcggtgc | |
| SJX1599 | tactgacctgcgaagcggctggacaggaa | |
| SJX1600 | tccagccgcttcgcaggtcagtatcatggccg | |
| SJX1601 | cgggccgtaccgcgcccagcggcgcgga | |
| SJX1602 | gctgggcgcggtacggcccgcagggcct | |
| SJX1603 | cggcgagatgacctcatgctgttttagagctagaaatagcaagttaaaataaggc | |
| SJX1604 | gcatgaggtcatctcgccgactagtattatacctaggactgagctagc | |
| SJX1727 | ggcttcaggcttcttcagcggttttagagctagaaatagcaagttaaaataaggc | pSJX046 |
| SJX1728 | cgctgaagaagcctgaagccactagtattatacctaggactgagctagc | |
| SJX1796 | ttgggcctggtgatcgacgcgttttagagctagaaatagcaagttaaaataaggctag | pSJX047 |
| SJX1797 | gcgtcgatcaccaggcccaaactagtattatacctaggactgagctagctg | |
| SJX1741 | gtcggtgctttttttcgtgcagtacatcgccacc | pSJX048 |
| SJX1746 | gcgatgtactgcacgaaaaaaagcaccgactcgg | |
| SJX1742 | ccagagcgccggccagtcgactttttagcgc | |
| SJX1743 | tcgactggccggcgctctggtcttaccg | |

| | | |
|---|---|---|
| SJX1744 | gctgggcgcggactcgtcaacatcccgcag | |
| SJX1745 | ttgacgagtccgcgcccagcggcgcgga | |
| SJX1764 | gctgagctgatccacgagaggttttagagctagaaatagcaagttaaaataaggctag | |
| SJX1765 | ctctcgtggatcagctcagcactagtattatacctaggactgagctagctg | |
| SJX1766 | ggtctggagaccttgctcgagttttagagctagaaatagcaagttaaaataaggctag | pSJX049 |
| SJX1767 | tcgagcaaggtctccagaccactagtattatacctaggactgagctagctg | |
| SJX1768 | accgccggagttcagctgcggttttagagctagaaatagcaagttaaaataaggctag | pSJX050 |
| SJX1769 | cgcagctgaactccggcggtactagtattatacctaggactgagctagctg | |
| SJX1613 | gtcggtgctttttttacctcgactatgagcggatcg | pSJX051 |
| SJX1614 | gctcatagtcgaggtaaaaaaagcaccgactcggtg | |
| SJX1615 | ttcgtccgtatcgggagatgcactcgacgcc | |
| SJX1616 | tcgagtgcatctcccgatacggacgaacacctgc | |
| SJX1617 | cgacgcagcgcgcgcccagcggcgcgga | |
| SJX1618 | gctgggcgcgcgctgcgtcgcggcgagg | |
| SJX1723 | gcgcaagtacgagaccgagagttttagagctagaaatagcaagttaaaataaggctagtc | |
| SJX1724 | tctcggtctcgtacttgcgcactagtattatacctaggactgagctagctg | |
| SJX1790 | gcatctcgaagacgtcgccggttttagagctagaaatagcaagttaaaataaggctag | pSJX052 |
| SJX1791 | cggcgacgtcttcgagatgcactagtattatacctaggactgagctagctg | |
| SJX1792 | atcctcgattgagagcgcaagttttagagctagaaatagcaagttaaaataaggctagtc | pSJX053 |
| SJX1793 | ttgcgctctcaatcgaggatactagtattatacctaggactgagctagctg | |
| SJX1605 | gtcggtgctttttttgttgcgggtgtactgctcgtgg | pSJX054 |
| SJX1606 | cagtacacccgcaacaaaaaaagcaccgactcggt | |
| SJX1607 | cccgacggagttctccgacgcgacgctga | |
| SJX1608 | gcgtcggagaactccgtcggggtcgcgga | |
| SJX1609 | gacgccccatcgcgcccagcggcgcgga | |
| SJX1610 | gctgggcgcgatggggcgtctgcgccgt | |
| SJX1717 | cgaaccccgaagatccaggggttttagagctagaaatagcaagttaaaataaggc | |
| SJX1718 | ccctggatcttcggggttcgactagtattatacctaggactgagctagc | |
| SJX1719 | tctcgttcagctgatcgacagttttagagctagaaatagcaagttaaaataaggc | pSJX055 |
| SJX1720 | tgtcgatcagctgaacgagaactagtattatacctaggactgagctagc | |
| SJX1798 | gctcgaagaagagctgaagggttttagagctagaaatagcaagttaaaataaggctag | pSJX056 |
| SJX1799 | ccttcagctcttcttcgagcactagtattatacctaggactgagctagctg | |
| SJX2395 | cccggctggatcaccggtttactagtattatacctaggactgagctagc | pSJX057 |
| SJX2396 | aaaccggtgatccagccggggttttagagctagaaatagcaagttaaaataaggc | |
| SJX2397 | gtgctttttttgccccaaaggcggccgaa | |
| SJX2398 | gcctttggggcaaaaaaagcaccgactcggtgc | |
| SJX2399 | ggaggccacccccctgtcggacgctgag | |
| SJX2400 | ccgacaggggggtggcctccgaagaagg | |
| SJX1595 | gctggaggggcgcgcccagcggcgcggatc | |
| SJX1596 | gctgggcgcgcccctccagccaggcgaaga | |
| SJX2401 | gctctattcctactgcaacggttttagagctagaaatagcaagttaaaataaggc | pSJX058 |
| SJX2402 | cgttgcagtaggaatagagcactagtattatacctaggactgagctagc | |
| SJX1784 | gttgggcgagaagaacgtcggttttagagctagaaatagcaagttaaaataaggctag | pSJX059 |
| SJX1785 | cgacgttcttctcgcccaacactagtattatacctaggactgagctagctg | |
| SJX1527 | tcggtgctttttttgacagcatgttggcgacg | pSJX060 |
| SJX1528 | cgccaacatgctgtcaaaaaaagcaccgactcggt | |
| SJX1529 | tcgggcgcactcttgagcccggccgccg | |
| SJX1530 | gggctcaagagtgcgcccgagcaatttaaag | |

| | | |
|---|---|---|
| SJX1576 | ccgcaaccgacgcgcccagcggcgcgga | |
| SJX1590 | gctgggcgcgtcggttgcggacgcaggc | |
| SJX1543 | gggtggcattaagagccgaggttttagagctagaaatagcaagttaaaataaggcta | |
| SJX1544 | ctcggctcttaatgccacccactagtattatacctaggactgagctagct | |
| SJX1545 | gatctggtcagcggtcaggggttttagagctagaaatagcaagttaaaataaggcta | pSJX061 |
| SJX1546 | ccctgaccgctgaccagatcactagtattatacctaggactgagctagct | |
| SJX1541 | cagggccccgatctgcactggttttagagctagaaatagcaagttaaaataaggcta | pSJX062 |
| SJX1542 | cagtgcagatcggggccctgactagtattatacctaggactgagctagctg | |
| SJX1876 | ggtcaatctaggcgatggtgagcaagggcgag | pSJX063 |
| SJX1877 | cccttgctcaccatcgcctagattgacccagtttgag | |
| SJX1878 | gagctgtacaagtaagtttcttgcctcgccacc | |
| SJX1879 | gagctgtacaagtaagtttcttgcctcgccacc | |
| SJX1876 | ggtcaatctaggcgatggtgagcaagggcgag | pSJX064 |
| SJX1877 | cccttgctcaccatcgcctagattgacccagtttgag | |
| SJX1878 | gagctgtacaagtaagtttcttgcctcgccacc | |
| SJX1879 | gagctgtacaagtaagtttcttgcctcgccacc | |
| SJX1968 | catcgaggcagaagatgcacgttttagagctagaaatagcaagttaaaataaggctagtc | pSJX065 |
| SJX1969 | gtgcatcttctgcctcgatgactagtattatacctaggactgagctagctg | |
| SJX1939 | cgcgcccagcggcgcggatccaaa | |
| SJX1940 | aaaaaaagcaccgactcggtgcca | |
| SJX1941 | tgatcggcacgtaagaggttc | |
| SJX1942 | ttacttgtacagctcgtccatgcc | |
| SJX1943 | accgagtcggtgcttttttccaaggcgcgctcccgggcgca | |
| SJX1944 | aacctcttacgtgccgatcatgtccgagcaggccgctctctg | |
| SJX1945 | tggacgagctgtacaagtaagattggagagcgcccctcggtc | |
| SJX1946 | gatccgcgccgctgggcgcgggttctatggtgctcatagtttccaaat | |
| SJX1978 | ggtcaacgcagcgccgaccagttttagagctagaaatagcaagttaaaataaggctagtc | pSJX066 |
| SJX1979 | tggtcggcgctgcgttgaccactagtattatacctaggactgagctagctg | |
| SJX1939 | cgcgcccagcggcgcggatccaaa | |
| SJX1940 | aaaaaaagcaccgactcggtgcca | |
| SJX1941 | tgatcggcacgtaagaggttc | |
| SJX1942 | ttacttgtacagctcgtccatgcc | |
| SJX1951 | accgagtcggtgcttttttatcgaggccgcccatctgccca | |
| SJX1952 | aacctcttacgtgccgatcacgctcgtcccaacaagagcc | |
| SJX1953 | tggacgagctgtacaagtaaagcgcccatgcaccggaggg | |
| SJX1954 | gatccgcgccgctgggcgcggcaacgtccaggagcagaactg | |
| SJX2018 | gaaggcgcccagatcgaagtgttttagagctagaaatagcaagttaaaataaggctag | |
| SJX2019 | acttcgatctgggcgccttcactagtattatacctaggactgagctagctg | pSJX067 |
| SJX1958 | accgagtcggtgcttttttgaaggtgggaatggtctgcatca | |
| SJX1940 | aaaaaaagcaccgactcggtgcca | |
| SJX1959 | gacgcagcggggatccactcccggagcgc | |
| SJX1960 | ggagtggatccccgctgcgtccggttgc | |
| SJX1961 | gatccgcgccgctgggcgcgatagagtaagccggccggacagcg | |
| SJX1939 | cgcgcccagcggcgcggatccaaa | |
| SJX2262 | cagatcgaagttggcggcaagttttagagctagaaatagcaagttaaaataaggctag | pSJX068 |
| SJX2263 | ttgccgccaacttcgatctgactagtattatacctaggactgagctagctg | |
| SJX2264 | agatcgaagttggcggcaacgttttagagctagaaatagcaagttaaaataaggctag | pSJX069 |
| SJX2265 | gttgccgccaacttcgatctactagtattatacctaggactgagctagctg | |

| | | |
|---|---|---|
| SJX2074 | ctatgacatgattacatggacaagaagtactcgatcgg | pSJX070 |
| SJX2075 | cgacggccagtgcctcatttgtacagttcatccataccatgcg | |
| SJX2076 | aactgtacaaatgaggcactggccgtcgtttt | |
| SJX2077 | gtacttcttgtccatgtaatcatgtcatagctgtttcctgtgtga | |
| SJX2078 | accgagtcggtgcttttttcgaagggcagttcgaccttg | |
| SJX2079 | ggcttgcgcggcccgctacatgacgggg | |
| SJX2080 | tgtagcgggccgcgcaagccgaccggcg | |
| SJX2086 | tcctttttaacccatttgacagctagctcagtcctagg | |
| SJX2087 | ttttcttttgcgtttggcgaaatcgtcaaaacggcttg | |
| SJX2089 | atgggttaaaaaggatcgatcctctagc | |
| SJX2142 | agctatgcgacgatgaatgcgttttagagctagaaatagcaagttaaaataaggctagt | |
| SJX2143 | gcattcatcgtcgcatagctactagtattatacctaggactgagctagctg | |
| SJX1940 | aaaaaaagcaccgactcggtgcca | |
| SJX2311 | aaacgcaaaagaaaatgccgattatgg | |
| SJX2144 | ggtcgcatcgcgtctcgcatgttttagagctagaaatagcaagttaaaataaggctagt | pSJX071 |
| SJX2145 | atgcgagacgcgatgcgaccactagtattatacctaggactgagctagctg | |
| SJX2146 | tctcctattgccgtcgccatgttttagagctagaaatagcaagttaaaataaggctagt | pSJX072 |
| SJX2147 | atggcgacggcaataggagaactagtattatacctaggactgagctagctg | |
